# B1 and B2 B cells are characterized by distinct CpG modification states at DNMT3A-maintained enhancers

**DOI:** 10.1101/2020.04.30.071209

**Authors:** Vinay S. Mahajan, Hamid Mattoo, Na Sun, Vinayak Viswanadham, Grace J. Yuen, Hugues Allard-Chamard, Maimuna Ahmad, Samuel JH Murphy, Annaiah Cariappa, Yesim Tuncay, Shiv Pillai

## Abstract

We show that DNA methylation is a layered process in B lymphocytes. An underlying foundational methylome is stably established during B lineage commitment and overlaid with a DNMT3A-maintained dynamic methylome which is sculpted in distinct ways in B1 and B2 B cells during B cell development. An engineered loss of DNMT3A after commitment to the B lineage unmasks a foundational methylome that is shared in both B1 and B2 sub-lineages. The dynamic methylome is comprised of novel enhancers whose methylation state is maintained by DNMT3A but can be modulated in strikingly different ways in B1 and B2 B cells. During B1 B cell development, the dynamic methylome undergoes a prominent programmed demethylation event that is not observed during B2 B cell development. The methylation pattern of the dynamic methylome is determined by the coincident recruitment of DNMT3A and TET enzymes and it regulates the developmental expression of B1 and B2 lineage-specific genes.

## Introduction

B cells are divided into two major sub-lineages. B2 B cells, which constitute the majority of all B cells, are short-lived recirculating cells that are metabolically quiescent and can be activated in secondary lymphoid organs where they collaborate with helper T cells to generate specific antibodies. B1a B cells, found predominantly in the peritoneal and pleural cavities, are long-lived self-renewing cells that produce natural antibodies as part of innate-like immunity ^1,2^. B1a B cells generally recognize ubiquitous multivalent antigens with relatively high affinity, supporting a role for BCR signaling in their generation or maintenance. There are two distinct models for B1a B cell development ^1^. The “lineage” model takes into account that these self-renewing B cells primarily arise from progenitors in the yolk sac and the fetal liver. B lineage progenitors derived from the fetal liver express the Lin28B RNA binding protein that represses the *let7* microRNA family and may consequently permit the expression of transcription factors such as Arid3a and Myc that help drive a proliferative program and B1a B cell development. Another key transcription factor that is subsequently required for B1a B cell development is Bhlhe41 ^1,3^. The “selection” model suggests that BCR signaling by ubiquitous antigens that bind to the BCR with relatively high affinity can induce the B1a B cell developmental program. Strong support for this model has been provided by the demonstration that replacing the BCR in B2 B cells with a B1a specific BCR can switch the B2 B cells to a B1a B cell fate ^4^.

Given the crucial role of DNA methylation in controlling gene expression, lineage specification, and the maintenance of cellular states, we explored the B1 and B2 sub-lineages through the lens of whole genome cytosine modification dynamics ^5,6^. Intergenic CpG island methylation has been linked to lineage specification in the immune system ^7^, and it has been suggested that DNA methylation might protect specific cell types from the aberrant activation of transcription factors in a lineage-specific manner ^8^. Genome-wide studies of cytosine methylation have been undertaken in human B cells, and it has been shown that early human B cell development is characterized by the loss of CpG methylation at enhancer sites; additionally, the differentiation of B cells into memory and plasma cells is characterized by the loss of CpG methylation in regions of heterochromatin and a gain of methylation in polycomb-repressed regions ^9^. However, the identification of human B1 B cells is still controversial and prior studies on human B cell methylomes did not address the B1 vs. B2 lineage decision. We have therefore explored this question in mice.

The enzymes that add methyl marks to cytosines include DNA methyltransferase 1 (DNMT1) and the DNA methyltransferases 3a (DNMT3A) and 3b (DNMT3B). DNMT1 is termed a maintenance methyltransferase because it recognizes hemi-methylated sites in newly replicated DNA and restores symmetric CpG methylation. DNMT3A and DNMT3B are *de novo* methyltransferases that are active during early embryogenesis and modify unmethylated CpGs symmetrically ^10^. Although 5-methylcytosine (5mC) is a stable modification, it can be converted to 5-hydroxymethylcytosine (5hmC) by methylcytosine dioxygenases of the Ten Eleven Translocation Family (TET1, TET2, and TET3) ^11–14^. 5hmC itself is also a substrate for TET enzymes and can be progressively oxidized to 5-formyl (5fC) and 5-carboxy (5caC) cytosine, which are recognized and excised by thymine-DNA glycosylase, such that an unmethylated cytosine is restored by the base-excision repair pathway. Unlike 5mC, the 5hmC mark is not maintained by DNMT1 during cell division and appears to undergo DNA replication dependent loss, thereby providing an additional mechanism for DNA demethylation ^15^. Thus, the TET enzymes play a critical role in active demethylation of mCpGs. Hypomethylated CpGs mark gene regulatory regions such as promoters and some enhancers. The 5hmC mark may serve as an indicator of TET activity, but given that this mark is especially enriched in enhancer elements, it has been speculated that it may play an active, functional role beyond being just an intermediate in demethylation ^16–18^. Several readers that specifically recognize methylated or unmethylated CpGs have been identified, but a specific reader for 5hmC has not yet been identified. Unlike DNMT1, which is associated with the DNA replication machinery, DNMT3A and the TET enzymes are recruited to specific genomic sites such as gene regulatory regions through their association with transcription factors. Thus, analysis of genome-wide CpG modifications can provide a window into the transcriptional regulatory landscape of a cell.

Once a CpG methylation pattern is established across the genome by DNMT3A and TET enzymes during B lineage commitment, it is subsequently maintained across cell divisions by DNMT1. Mice lacking *Tet2 and Tet3* in the B lineage exhibit normal numbers of B2 B cells but experience a partial block in early B cell development and completely lack B1a and marginal zone B cells ^19^, suggesting a continued role for DNA demethylation in establishing the B1a B cell program after B lineage commitment.

We used whole genome bisulfite sequencing to profile DNA methylation in B1 and B2 B cells and their precursors in mice. We also used a conditional allele of DNMT3A, which is deleted after B lineage commitment, and showed that both B1 and B2 B cells express a dynamic layer of B1 and B2 sub-lineage specific and DNMT3A-maintained CpG methylation that is superimposed on a shared foundational methylome. We show that continued action of DNMT3A is required to counteract TET activity in this dynamic methylome layer. The sites whose CpG methylation is maintained by DNMT3A after B lineage commitment are highly enriched for enhancers, that we have termed DNMT3A-maintained enhancers (DMEs). Our results, illustrating that B1 and B2 B cells share a nearly identical foundational methylome, but with superimposed developmentally regulated and dynamically modulated lineage-specific methylomes, provide an epigenetic framework for understanding the reported BCR-dependent plasticity in B cell sub-lineages.

We show here that programmed demethylation of a large set of lineage-specific transcriptional enhancers occurs in developing B1a B cells but not in B2 B cells, and that the demethylated enhancers are linked to genes that are part of the B1a B cell developmental program. We also show that DNMT3A maintains the CpG methylation of a novel subset of enhancers in a lineage-specific manner in B cells. Based on the distinctive properties of these enhancers, we refer to them as DNMT3A-maintained enhancers (DMEs). In B2 B cells, DMEs have the hallmarks of active enhancers, exhibit H3K27ac and H3K4me1 marks, lack the H3K4me3 modification, but also exhibit high levels of 5mCpG and 5hmCpG marks, indicative of concurrent DNMT3A and TET enzyme activity at these sites. In B1a B cells, these very same DMEs acquire non-canonical H3K4me3 marks during development, concurrent with the absence of both 5mCpG and 5hmCpG. The demethylation or methylation of DMEs is linked to the regulated expression of lineage-specific genes and has revealed a novel functional role for DNMT3A in both marking and maintaining DNA methylation at specific enhancers that modulate the fate of developing lymphocytes. This study therefore illustrates how DNMT3A functions as a maintenance methyltransferase across a broad swath of B lineage-specific enhancers in a manner that is distinct from the DNA replication-coupled maintenance methyltransferase function of DNMT1. Our results also provide additional insights into how perturbation of the dynamic methylome promotes the pre-leukemic phase of IgHV-unmutated chronic lymphocytic leukemia, a leukemia suspected to be of B1 B cell origin.

## Results

### B1a B cell development is characterized by a pronounced loss of CpG modification

We surveyed the genome-wide CpG modification profiles in murine B1a and B2 B cell lineages, by performing whole genome bisulfite sequencing (WGBS) of proB2 cells (Hardy fractions B and C from bone marrow), splenic follicular B2 cells, proB1 cells (Hardy fractions B and C from fetal liver) and peritoneal B1a B cells; these populations will hereon be referred to as proB2, B2, proB1 and B1 B cells, respectively. We identified about 40,000-50,000 hypomethylated regions (HMRs) that were dispersed across 1.7% to 1.9% of the genome, largely in intergenic and intronic regions in all four cell types ^20^. Interestingly, the HMRs in wild-type B1 B cells were found to be significantly longer than those in the B2 B cells (median lengths of 830 bp and 668 bp, respectively; K-S test p << 1e-10), indicating that the B1 lineage may favor CpG demethylation at HMRs across the genome **(Figure 1A)**. Large HMRs over 3.5 kb have been previously referred to as canyons in studies on murine hematopoietic stem cells ^21^. Using a similar cutoff of 3.5 kb, we identified methylation canyons in all four B cell populations. Mature B1 B cells exhibited 30-35% more canyons than all other B cell stages examined **(Figure 1B)**. This developmental increase in the number of canyons in B1 B cells was reminiscent of the canyon expansion observed in *Dnmt3a*-deficient murine HSCs ^21^ and suggested that changes in DNMT3A activity may contribute to the programmed demethylation observed during B1 B cell differentiation.

**Figure 1:**
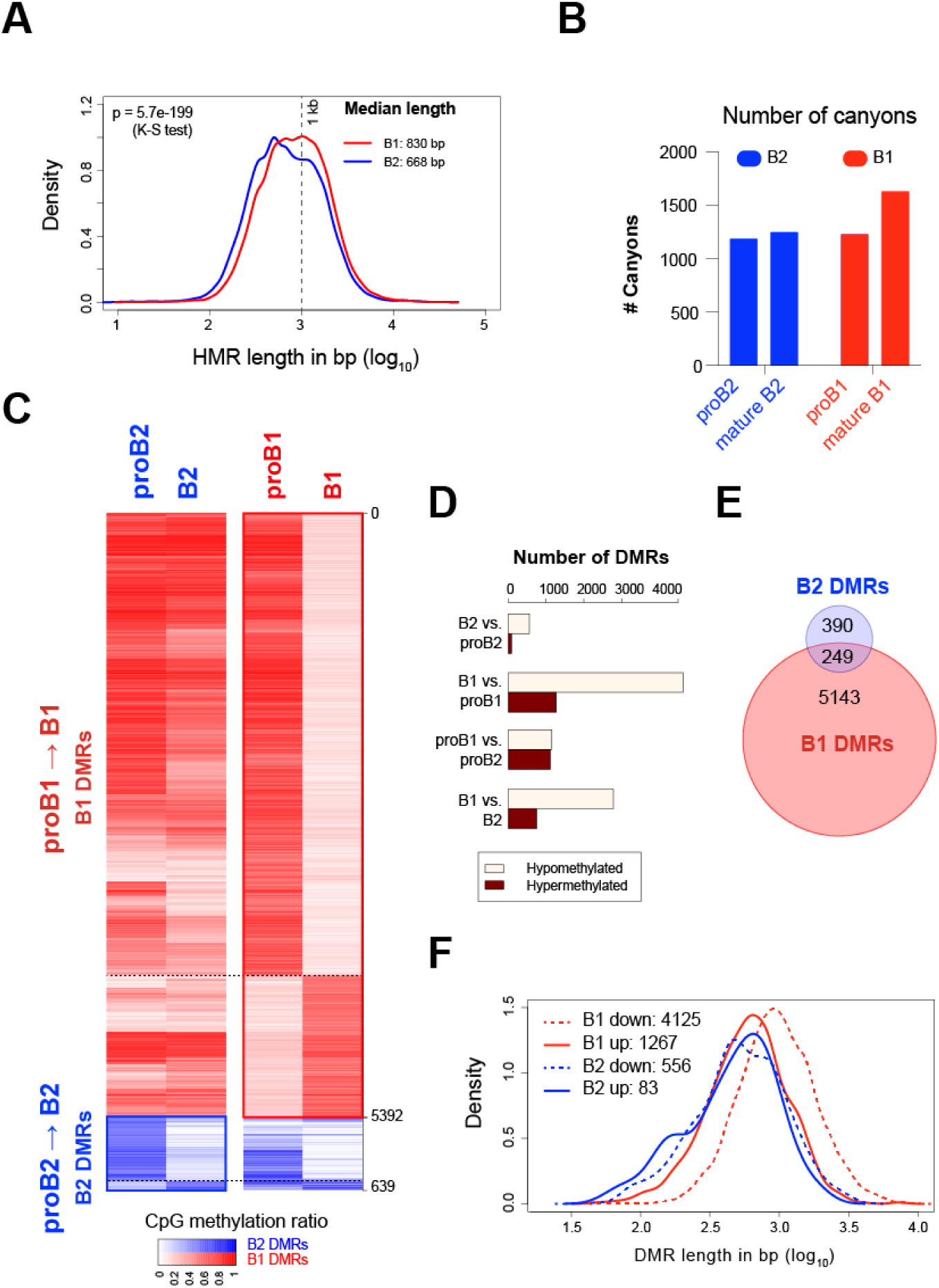
B1a B cell development is marked by a pronounced loss of CpG modification. (A) Distribution of HMR lengths in proB and B cells in the B1 and B2 lineage. (B) Number of canyons (HMR length > 3.5 kb) identified in proB2, B2 (follicular B), proB1 and B1 B cells. (C) CpG modification profiles in developmental DMRs in the B1 and B2 lineages. Average CpG modification ratio in the B1 and B2 DMRs in proB1, B1, proB2 and B2 B cells is depicted in the heatmap. (D) The number of DMRs identified by pairwise comparisons between proB and mature B cells and between corresponding stages of the B1 and B2 B cell lineages. (E) Overlap between B2 and B1 DMRs; DMRs with over 50% overlap were considered overlapping. (F) Distribution of development DMR lengths in proB and B cells in the B1 and B2 lineage.

We next identified differentially methylated regions (DMRs) that arise during the development of the B1 and B2 B cell lineages. DMRs are regions with a high probability of differential methylation where an HMR exists in one methylome but not at the same location in the other. DMRs were particularly abundant between the proB1 and B1 stages. They encompassed approximately 6 million bp across 5,392 genomic intervals in the B1 lineage (0.42% of the genome), but only about 0.5 million bp across 639 intervals in the B2 lineage (0.04% of the genome), suggesting that the regulation of CpG methylation is a far more prominent feature of B1a B cell development, compared to B2 B cell development **(Figure 1C-E)**. The majority of the B1 DMRs (∼75%) exhibited a loss of CpG modification during B1 B cell development (**Figure 1C, D).** The median size of the observed B1 DMRs was 857 bp, and approximately 12% of the hypomethylated B1 DMRs fell within methylation canyons in B1 B cells (**Figure 1F**)

### Programmed demethylation in B1a B cells occurs at enhancers that are demethylated and re-methylated in B2 B cells

The majority of B1 and B2 DMRs were present at a distance from the transcription start site (TSS) in intergenic or intragenic spaces, compatible with a possible enhancer function **(Figure 2A, B)**. Indeed, many prior studies of CpG methylomes in adult somatic tissues have found that variably methylated regions are strongly enriched at enhancer sites. Analysis of publicly available ChIP-Seq data for H3K4me1 and H3K4me3 modifications in mature B2 and B1 B cells ^22^, showed that the DMRs that underwent developmental demethylation in both B1 and B2 B cells co-localized with H3K4me1 peaks in mature B1 and B2 B cells, suggesting that these DMRs might represent developmentally demethylated enhancers that modulate lineage-specific gene expression **(Figure 2C)**. To assess the significance of these enhancers in the context of hematopoietic development, we also examined H3K4me1 and H3K4me3 marks in HSCs using published ChIP-Seq data. The DMRs that were developmentally hypermethylated in mature B2 or B1a B cells exhibited little H3K4me1 signal in B cells, but did bear an H3K4me1 signature in HSCs, suggesting that these hypermethylated DMRs may represent enhancers that were developmentally silenced by CpG methylation **(Figure 2C)**. The B1 DMRs exhibited low levels of H3K4me1 in HSCs suggesting that they may represent primed enhancers in HSCs that subsequently become fully active in the B lineage.

**Figure 2:**
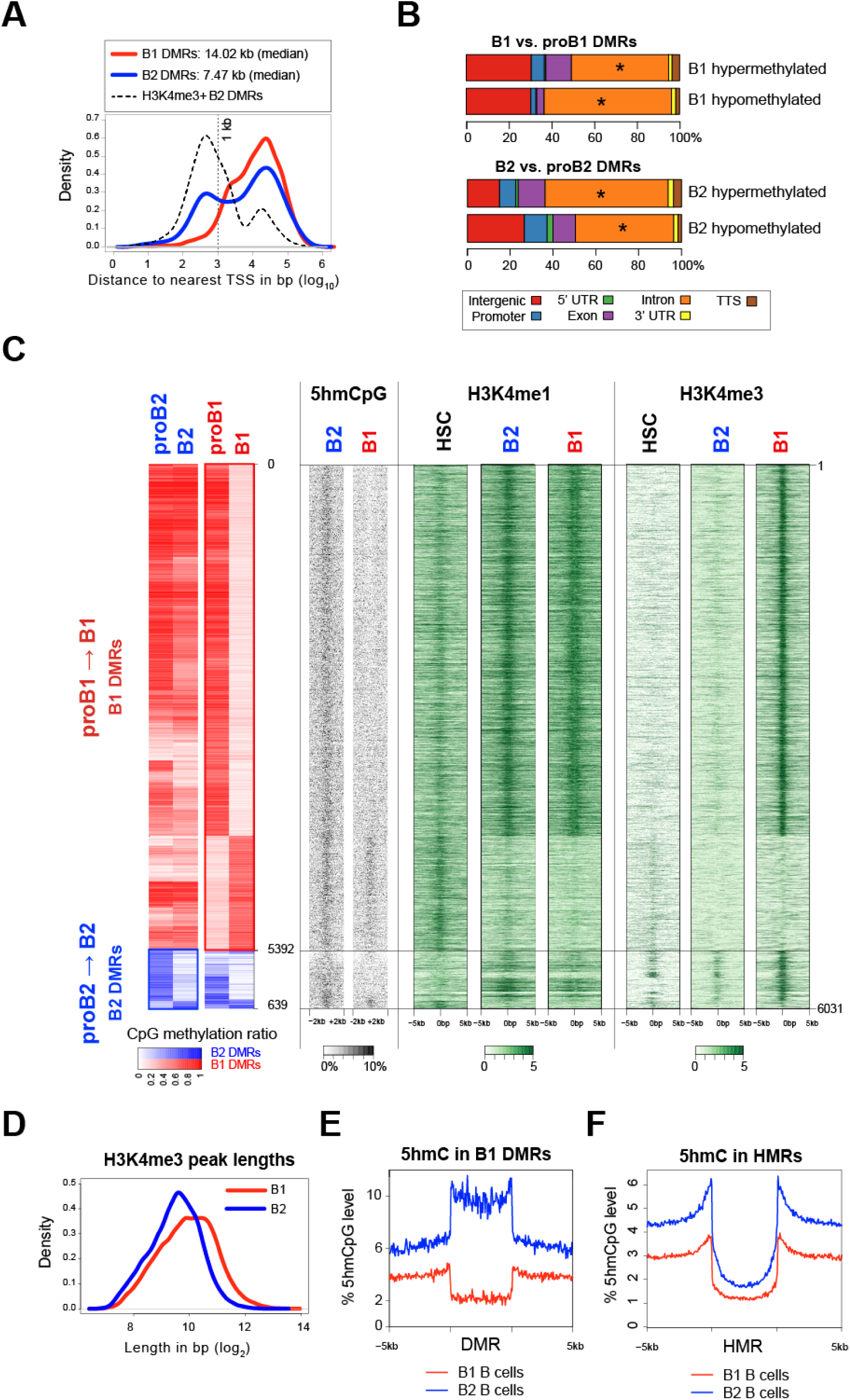
Extensive loss of enhancer methylation during B1a B cell development. (A) Distribution of distance to nearest TSS for B1 (red) and B2 (blue) DMRs. B2 DMRs that overlap with H3K4me3 ChIP-Seq peaks in B2 B cells (black) are separately depicted. (B) DMR genomic features. Asterisks depict genomic features that are significantly enriched with respect to HMRs. (C) 5hmCpG, H3K4me1 and H3K4me3 histone marks in and around the corresponding B1 and B2 DMRs in HSCs ^21^, B1 and B2 B cells ^22^ are depicted as a heatmap. (D) Distribution of H3K4me3 peak lengths in B1 (red) and B2 (blue) cells. (E) Average profile of 5hmCpG modification (% of total CpGs) in the B1 DMRs in B1 (red) and B2 (blue) B cells. The DMRs are scaled to the same length and 5 kb flanking regions are depicted. (F) Average profile of 5hmCpG modification (% of total CpGs) in the HMRs in B1 and B2 B cells and 5 kb flanking regions.

In B2 B cells, only a small subset of developmentally demethylated DMRs that were present in the close proximity of annotated transcription start sites (TSS) exhibited an H3K4me3 signal, consistent with a promoter function **(Figure 2A)**. However, in B1 B cells, most DMRs that were developmentally demethylated acquired some H3K4me3 trimethylation marks **(Figure 2C)**. Although H3K4me3 is generally associated with promoters, it can be found in enhancers in some contexts ^23,24^. The recognition of unmethylated CpGs by the SET1 H3K4 trimethylase complex through CFP1 has been reported to restrict this chromatin modifying enzyme complex to unmethylated promoter regions and prevent CpG methylated enhancers from acquiring ectopic H3K4me3 marks ^25^. This CpG methylation barrier to H3K4me3 acquisition appears to break down at many enhancer sites in B1 B cells, and the developmental acquisition of non-canonical or ectopic H3K4me3 marks at demethylated enhancers is unique to B1 B cells. Interestingly the H3K4me3 peaks were wider in B1 B cells than B2 B cells across the genome, consistent with a global increase in HMR lengths in B1 B cells **(Figure 2D)**.

WGBS revealed that 90% of the CpGs in DMRs that were developmentally demethylated in B1 B cells were modified in B2 B cells (**Fig 2C**). As WGBS does not distinguish between cytosine methylation and hydroxymethylation, we also analyzed 5-hydroxymethylation (5hmC) levels in mature B1 and B2 B cells using TET-assisted bisulfite sequencing (TAB-Seq) ^26^ (**Fig 2C**). Given that the bulk of the bisulfite protection arose from 5mCpG rather than 5hmCpG, we focused on 5mCpG in our analyses. The majority of CpGs in B1 DMRs were modified in B2 B cells. About 10 percent of these CpG were hydroxymethylated, indicating that TET enzymes are active at these sites even in B2 B cells **(Figure 2E)**. In contrast, our analyses showed no appreciable level of 5hmC at the B1 DMRs in B1 B cells. As previously described in HSCs, 5hmCpGs, a mark of TET activity, were found to be enriched at HMR boundaries in both B1 and B2 B cells **(Figure 2F)**. TET activity at HMR boundaries plays a critical role in maintaining the HMRs by opposing the action of DNMT3A ^21,27^.

We conjectured that these findings may be best explained by a global reduction in DNMT3A activity during B1 B cell development, resulting in unopposed TET activity both at HMR boundaries as well as at B1 DMRs. This results in longer HMRs and increased hypomethylation canyons during B1 development, a phenotype previously linked to *Dnmt3a* deficiency in HSCs. Furthermore, unopposed TET activity at B1 DMRs in B1 B cells may result in more complete oxidation of 5hmC to 5-formyl or 5-carboxyl cytosines, which are rapidly recognized and replaced with unmodified cytosines by base excision repair.

### B1 lineage-specific gene expression is linked to enhancer demethylation

Examination of gene expression data in B cell subsets from the ImmGen Consortium indicated that the genes in the proximity of B1 B cell specific DMRs that lose methylation during B1 B cell development are upregulated during differentiation from proB1 to mature B1 B cells (either fetal liver Fraction E or peritoneal B1a B cells) **(Figure 3A)** ^28^. These genes are enriched for pathways involved in B cell receptor signaling and activation **(Figure 3B)**. In contrast, B1 B cell-specific DMRs that gain methylation during development were not associated with statistically significant changes in the expression of neighboring genes **(Figure 3A)**. B2 DMRs were less numerous and exhibited little correlation with gene expression changes during development **(Figure S2)**. Genes whose expression was increased during B1 B cell development exhibited a significant overlap with genes that were closest to DMRs that lost CpG methylation during B1 B cell development (p < 0.01; Fisher’s exact test) **(Figure 3C)**. Of these genes, 40% encode B1 lineage-specific genes, that include transcription factors such as *Arid3a, Bhlhe41* and *Zbtb32*, and signaling regulators such as *Siglecg* and *Rasgrp1*, that are known to be required for B1 B cell development and function (**Figure 3D)** ^29–32^.

**Figure 3:**
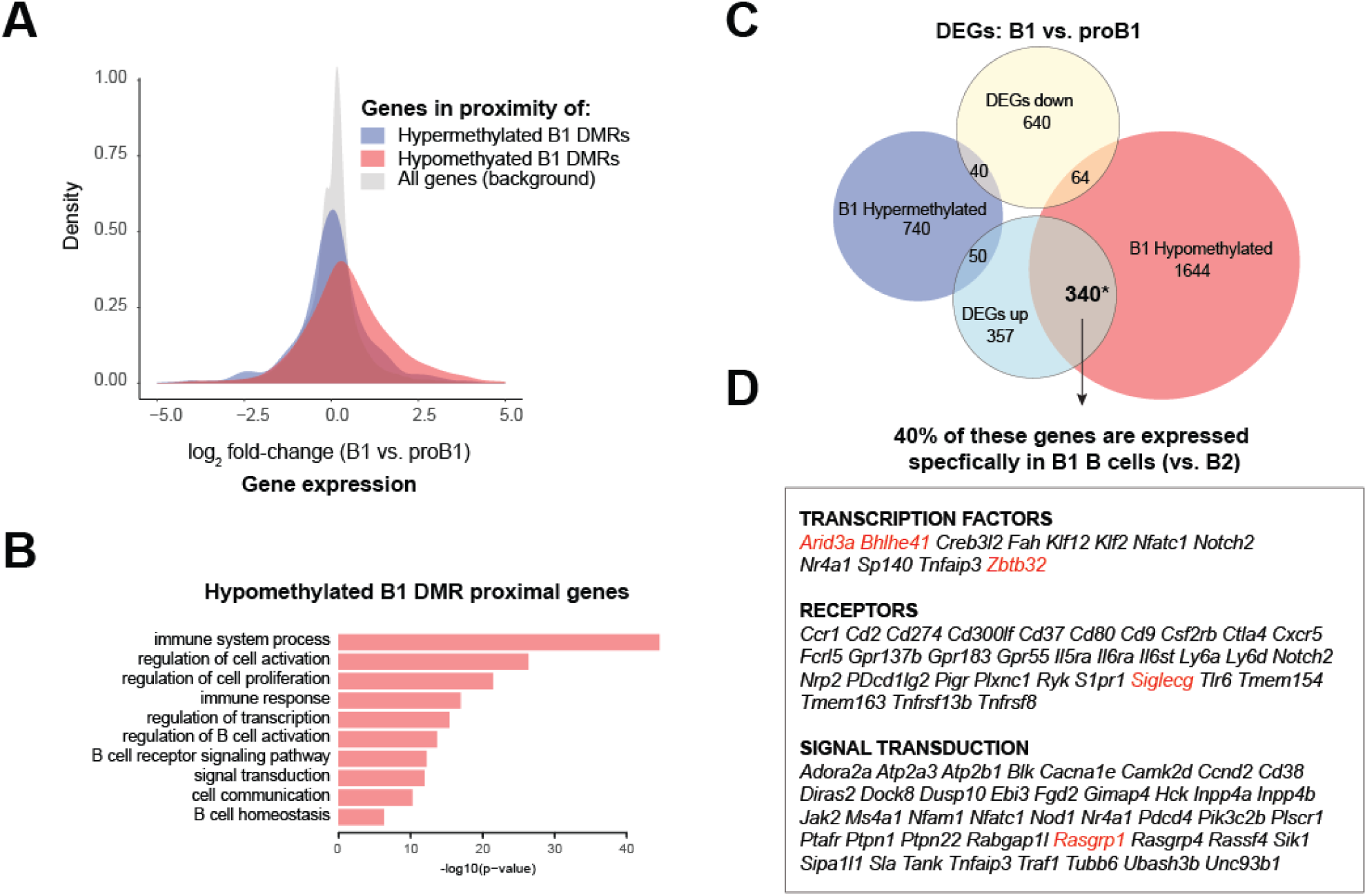
B1 lineage-specific gene expression is linked to enhancer demethylation. (A) Distribution of fold-changes in gene expression between mature B1 (peritoneal B1a or fetal liver FrE) and proB1 (fetal liver FrBC) cells. Genes in the proximity of developmentally hypomethylated or hypermethylated B1 DMRs are depicted alongside the set of all genes. Microarray data from the ImmGen Consortium was used for this analysis. (B) Enriched Gene Ontology terms for genes in the proximity of hypomethylated B1 DMRs. (C) Overlap between differentially expressed genes and genes in the proximity of DMRs between B1 B (peritoneal B1a or fetal liver FrE) and proB1 (fetal liver FrBC) cells. Gene sets with a significant overlap are marked with an asterisk (p < 0.001, Fisher’s exact test). (D) B1 lineage-specific genes in the significantly overlapping set shown in (C).

### Deficiency of Dnmt3a in the murine B lineage results in the selective expansion of B1a B cells

In order to evaluate the possible contribution of DNMT3A to the B1 vs. B2 B cell fate decision, we used *Cd19-Cre* to conditionally delete a floxed *Dnmt3a* allele (*Dnmt3a*^*2lox*^) in the B lineage, specifically from the proB1 and proB2 stages onwards ^33,34^.The direct or indirect loss of DNMT3A in the B lineage has been shown to result in the leukemic transformation of B1 B cells^35–37^. Similarly, *Cd19-Cre*^*+/−*^ *Dnmt3a*^*2lox/2lox*^ (*Dnmt3a*^*−/−*^) mice also exhibited a marked expansion of B1a B cells that progressed to monoclonal leukemia by 9 months of age with a 100% penetrance **(Figure 4A)**. The increase in B1 B cells was initially detectable in the peritoneal cavity as early as 4 weeks of age **(Figure 4A)** and became evident in the spleen and bone marrow after 10 weeks of age. The peritoneal B1b B cell population was progressively displaced by the expanding B1a population with age. We observed no increase in the proportion of CD19^+^ B1a lineage progenitors or proB1 cells in the fetal liver ^38^, suggesting that the expansion of B1a B cells was driven by increased self-renewal or proliferation in the mature B1a B cell compartment **(Figure S3A)**. There was a slight decrease in the number of follicular B cells in the spleen and no change in the number of developing B2 B cells in the bone marrows of *Dnmt3a*^*−/−*^ mice **(Figure 4B and S3B)**. The number of marginal zone (MZ) B cells was also reduced at 8 weeks of age **(Figure 3B and S3C)**. Changes in the MZ B cell compartment could not be assessed beyond 12 weeks of age, as the splenic marginal zone niche was obliterated by infiltrating B1a cells and MZ B cells were no longer detectable (data not shown).

**Figure 4:**
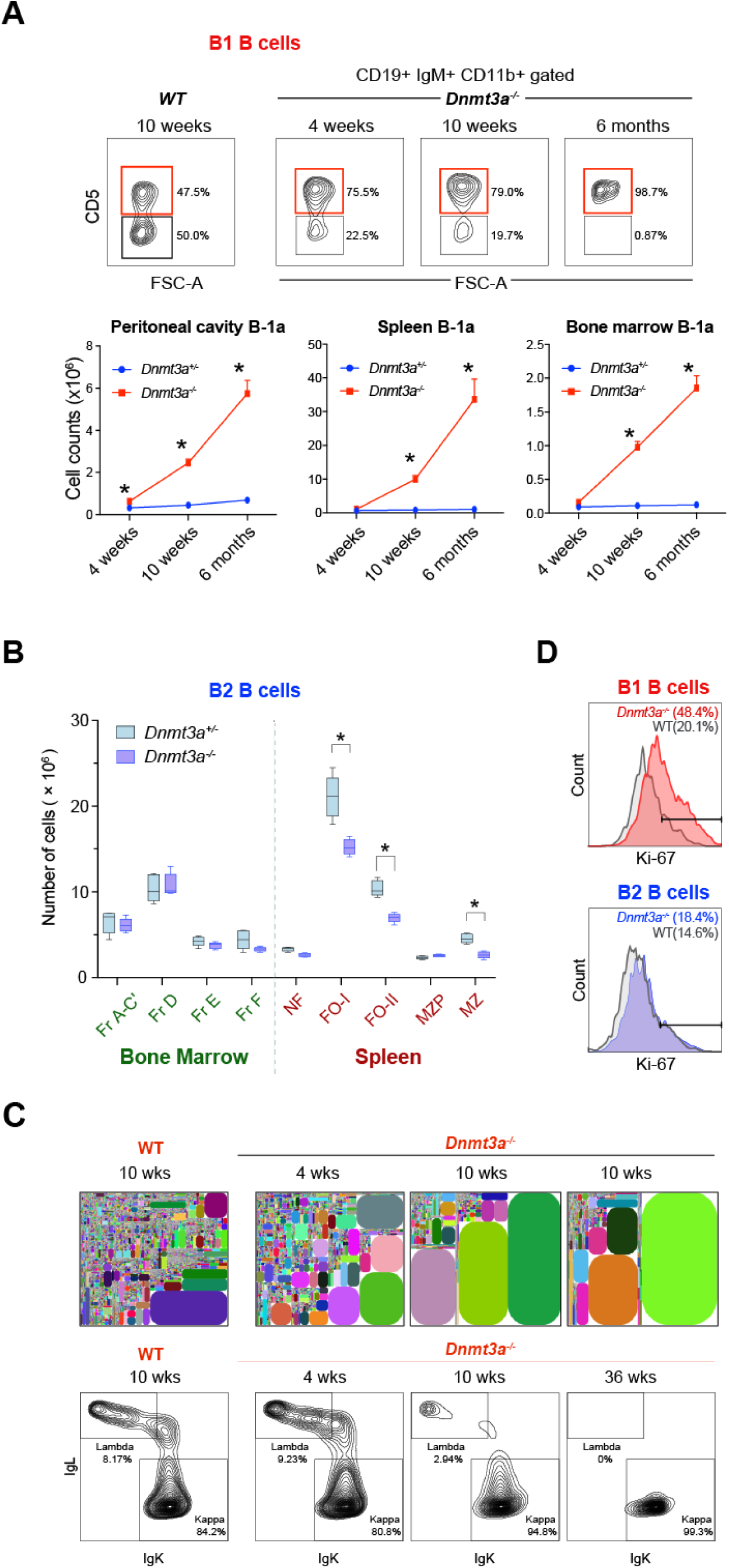
*Dnmt3a* deficiency in the B lineage results in the selective expansion of B1 B cells. (A) Expansion of B1a B cells (CD19^+^ IgM^+^ CD11b^+^ CD5^+^) in the peritoneal cavity, spleen and bone marrow of *Dnmt3a*^*−/−*^ mice. Five wild-type and *Dnmt3a*^*−/−*^ mice were analyzed in each group. Statistically significant differences are indicated with asterisks. (B) Number of bone marrow B2 B cell precursors (Fractions A-C’, D, E, F) and mature splenic B2 B cells (newly formed (NF), follicular (FO-I and FO-II), marginal zone precursor (MZP), and marginal zone (MZ) B cells) from 8 week old *Dnmt3a*^*−/−*^ mice and their littermate controls. The experiment was performed two times and a total of 8 mice in each group were analyzed. Plots represent mean ± SEM. Statistically significant differences (p < 0.05) are indicated with asterisks. (C) IgH repertoire of peritoneal B1a B cells in wild-type and *Dnmt3a*^*−/−*^ mice at varying ages (top row). Each clonotype is represented by a distinctly colored rectangle, with an area proportional to the fraction of total reads. Kappa and lambda Ig light chain staining in B1a B cells from wild-type and *Dnmt3a*^*−/−*^ mice at varying ages is also shown (bottom row). (D) Ki-67 levels in B1a B cells from the peritoneal cavity and B2 B cells from the spleens of 4 week old wild-type and *Dnmt3a*^*−/−*^ mice.

At 4 weeks of age, B1a B cells lacking DNMT3A remained largely polyclonal, similar to wild type B1a B cells, but by 10 weeks, they were significantly oligoclonal. Staining of Igκ and Igλ light chains showed that the expanded B1a B cell pool in the peritoneal cavity underwent a gradual restriction in the repertoire from a polyclonal to monoclonal state **(Figure 4C)**. Of the eight mice analyzed beyond 30 weeks of age, seven showed only Igκ staining while one showed only Igλ staining. *Dnmt3a*^*−/−*^ B1a B cells exhibited higher Ki-67 staining, consistent with increased self-renewal in this compartment **(Figure 4D)**. IgH rearrangements from the clonal leukemias were amplified and sequenced, and like unmutated human CLL, were found to be free of somatic hypermutation. To avoid potential confounding effects arising from oncogenic somatic mutations that may have been selected in a leukemic *Dnmt3a*^*-*/-^ clone at a later time point, we used peritoneal *Dnmt3a*^*-*/-^ B1a B cells from 4 week-old mice, when the cells were still in a polyclonal state, for all subsequent analyses.

### DNMT3A loss reveals a B lineage-specific foundational methylome shared in B1 and B2 B cells

In the absence of DNMT3A, lineage specific methylation patterns in mature B1 and B2 B cells were almost totally erased, unmasking a foundational methylome that was virtually identical in B1 and B2 B cells. Indeed, there were only 34 differentially methylated intervals between *Dnmt3a*^*−/−*^ B1 and B2 B cells **(Figure 5A, B)**. In contrast, we identified a total of 9,401 and 9,765 genomic intervals in the B2 and B1 B cell lineages, respectively, that exhibit differential CpG modification across proB, mature B (wild-type) and *Dnmt3a*-deficient mature B cells, and henceforth refer to them as DNMT3A-dependent DMRs (DDMRs). We found that the number of canyons increased in *Dnmt3a*-deficient B cells compared to their wild-type counterparts **(Figure 5C)** and appeared to arise from the expansion and amalgamation of smaller HMRs present at the respective sites in wild-type B cells. The canyons in *Dnmt3a*-deficient B2 and B1 B cells exhibited a very high degree of overlap (98%) and were strongly enriched for genes encoding transcription factors, such that for nearly one-third of these canyons, the closest gene encoded a transcription factor. Interestingly, the CpG methylation profile of wild-type B1a B cells resembled *Dnmt3a*^−/−^ B1 and B2 B cells, suggesting that DNMT3A activity may be physiologically downregulated during B1 B cell development **(Figure 5A, B).**

**Figure 5:**
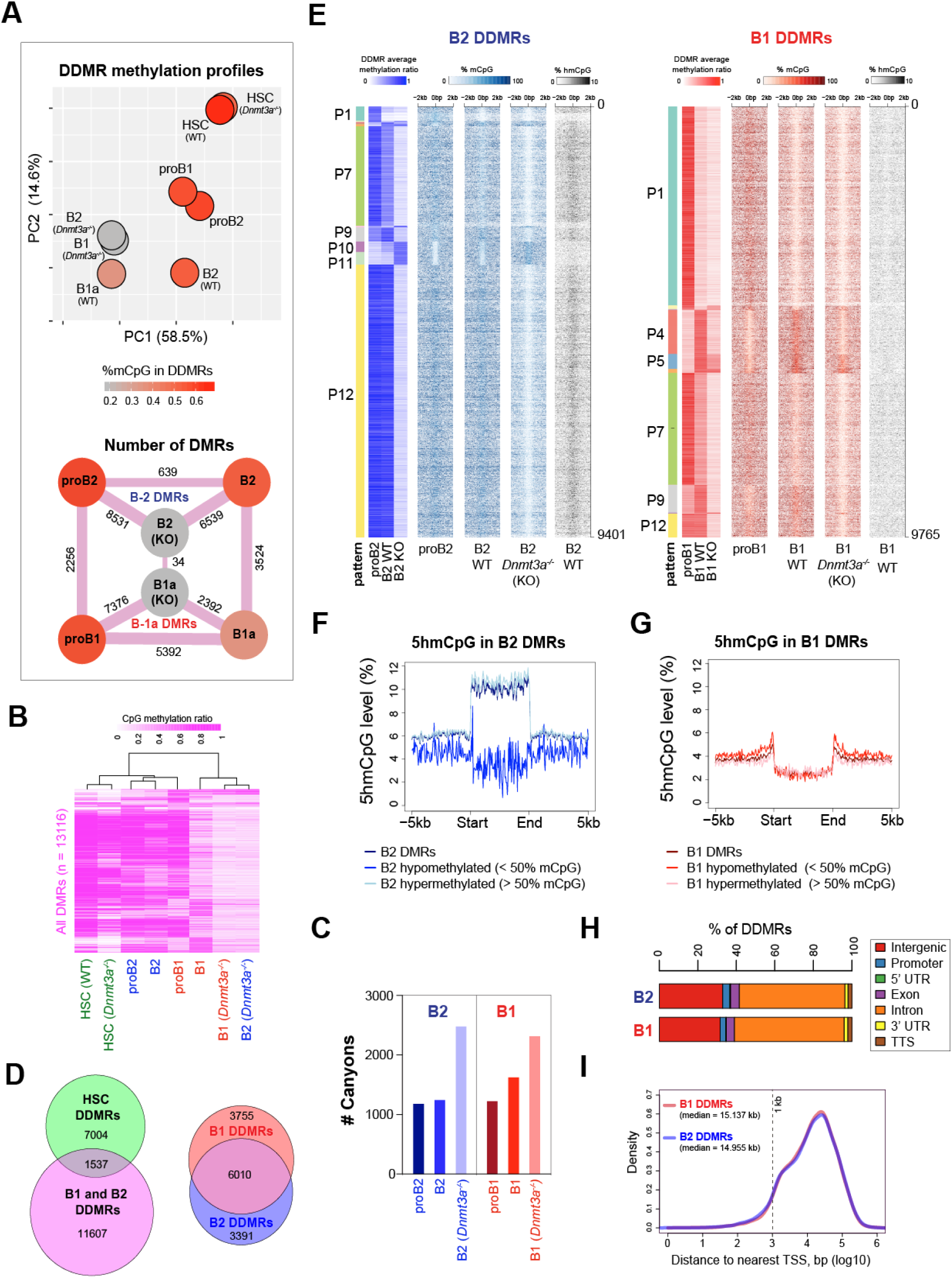
DNMT3A maintains lineage-specific CpG methylation patterns in B1 and B2 B cells. (A) Differences in the methylation profiles in the B lineage DDMRs (DNMT3A-dependent DMRs) observed across proB, mature B and *Dnmt3a*^*−/−*^ B cells in B2 and B1a B cell lineages as well as wild-type and *Dnmt3a*^−/−^ HSCs are depicted as a PCA plot (top) and as a graph of the number of the pair-wise DMRs (bottom). The %CpG methylation in the DDMRs for each condition is depicted using a color scale. (B) Differences between the CpG methylation profiles in the B lineage DDMRs across the eight conditions depicted in (A) are illustrated as a hierarchically clustered heatmap. (C) Number of canyons (HMR length > 3.5 kb) observed in proB, wild-type mature B and *Dnmt3a*^*−/−*^ B cells in the B2 and B1a lineages. (D) Degree of overlap between the B lineage DDMRs and HSC DDMRs (wild-type vs. *Dnmt3a*^*−/−*^ HSCs) as well as B1 and B2 DDMRs. (E) DNMT3A-dependent differentially methylated regions (DDMRs) in the B2 (n=9,401) and B1 (n=9,765) B cell lineage. Heatmaps showing the average methylation ratio within the B1a and B2 DMRs as well as the %CpG modification in 100 bp bins depicted ±2 kb from the center of the DMRs are shown for wild-type proB2, B2, proB1, B1 as well as *Dnmt3a*^*−/−*^ B1 and B2 B cells. The CpG methylation state of B2 DMRs is shown in shades of blue and B1 DMRs is shown in shades of red. The six main patterns of methylation observed for B1 and B2 B cells are labeled (P1-P12). (F and G) Average 5hmCpG modification level (% of total CpG) in B2 DDMRs in B2 cells (F) and B1 DDMRs in B1 cells (G). The DDMRs are scaled to the same length and depicted alongside 5 kb flanking regions. (H) Genomic features of DDMRs in B2 and B1 B cells. (I) Distribution of distances to the nearest TSS for B1 and B2 DDMRs.

Comparison with publicly available WGBS data from murine bone marrow hematopoietic stem cells (HSCs) suggested that B-lineage DDMRs were predominantly inherited in a methylated state from HSCs **(Figure 5A, 5B)** ^21^. These B lineage DDMRs were predominantly methylated in HSCs, proB1 and proB2 cells and their methylation state was broadly similar across these cell types. Given the use of *Cd19-Cre* for deleting the *Dnmt3a* floxxed allele, the loss of DNMT3A in our studies occurred after B lineage commitment, and the HSCs in the *Cd19-Cre*^+^ *Dnmt3a*^*−/−*^ mice were DNMT3A-sufficient. Interestingly, over three-fourths of the B lineage DDMRs retained their CpG methylation in HSCs following the loss of DNMT3A in those cells **(Figure 5B and 5D)**. This suggests that the foundational methylome is recast during B lineage commitment. Indeed, the majority of the B lineage DDMRs do not require DNMT3A for the maintenance of their CpG methylation in HSCs but become dependent only upon differentiation into B cells, perhaps to counteract TET2-mediated demethylation as suggested by the enrichment of 5hmC at these sites **(Figure 5E, 5F, 5G)**. In this context, DNMT3A appears to function as a maintenance methyltransferase at the DDMRs even in non-dividing B2 B cells, in a manner that is distinct from the replication-coupled maintenance methylation function performed by DNMT1. The set of DDMRs may therefore be viewed as the DNMT3A-maintained dynamic methylome in B1 and B2 B cells.

The DDMRs were stratified into 12 groups based on their patterns of CpG modification **(Figure S4A)**; six patterns accounted for >97% of all DDMRs in B2 and B1 B cells **(Figure 5E)**. As previously observed, the majority (∼75%) of the B1 DDMRs exhibit a loss of CpG modification during the differentiation of proB1 cells into B1a B cells **(Figure 5E)**. Notably, the DDMRs that undergo partial demethylation from the proB1 to the mature B1a stage exhibited almost complete demethylation in *Dnmt3a*^*−/−*^ B cells (Pattern 7 in **Figure 5E**), suggesting that even the reduced level of CpG methylation observed in the DDMRs in wild-type B1a B cells is maintained by DNMT3A. In B2 B cells, which exhibited >65% CpG modification in the DDMRs, about 10% of the CpGs in the B2 DDMRs were 5-hydroxymethylated indicative of ongoing TET activity at these sites **(Figure 5F).** In contrast, our analysis showed no appreciable level of 5hmCpG at the B1 DDMRs in B1 B cells **(Figure 5G)**. This suggests that the balance between TET and DNMT3A activities is flipped in B1 and B2 B cells - while TET activity is adequately balanced by DNMT3A in B2 B cells, DNMT3A activity is physiologically dampened at these sites during B1 B cell development.

### DNMT3A-maintained enhancers (DMEs) represent a novel methylation-sensitive subset of enhancers

The DDMRs were predominantly in intergenic and intronic regions at a distance from the TSS, consistent with an enhancer function **(Figure 5H and 5I)**. To assess this, we examined chromatin features at the DDMRs using CUT&RUN with antibodies against H3K4me1, H3K4me3 and H3K27ac ^39^ and ATAC-Seq ^40^. In both B1a and B2 B cells, the DDMRs were enriched for the H3K4me1 histone modification, a widely studied mark of enhancers **(Figure 6A and 6B)**. Indeed, the DDMRs accounted for approximately 11% of all enhancers defined by non-TSS H3K4me1 peaks (7990 of 74344) in B cells. Developmentally hypomethylated DDMRs (B1 DDMR patterns 1 and 7, and B2 DDMR pattern 7), acquired H3K27ac marks, which is indicative of an active enhancer state, during development **(Figure 6A and 6B)**. Both B2 and B1 DDMRs overlapped with sites of enhanced chromatin accessibility, and this was inversely correlated with CpG methylation (Spearman’s correlation coefficient = -0.26 and -0.57, for B2 and B1a DDMRs respectively). Given that the majority of the DDMRs were partially hypomethylated in wild-type B1a B cells, the loss of DNMT3A had a more profound impact on the CpG methylation of DDMRs in B2 B cells, concomitant with a gain in chromatin accessibility **(Figure 6B and 6C)**. This suggests that CpG methylation may constrain chromatin accessibility at these DNMT3A-maintained enhancers that we refer to as DMEs.

**Figure 6:**
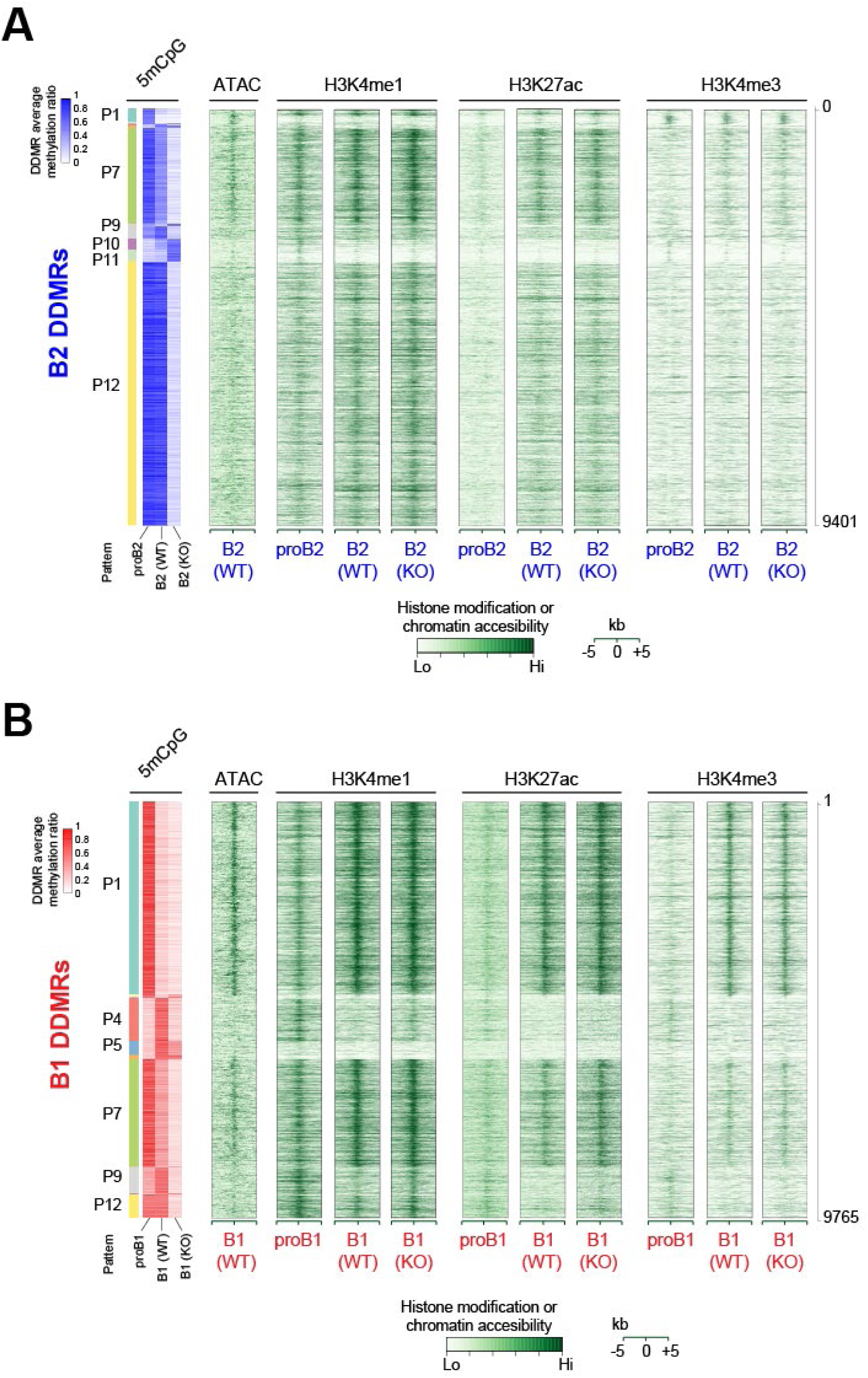
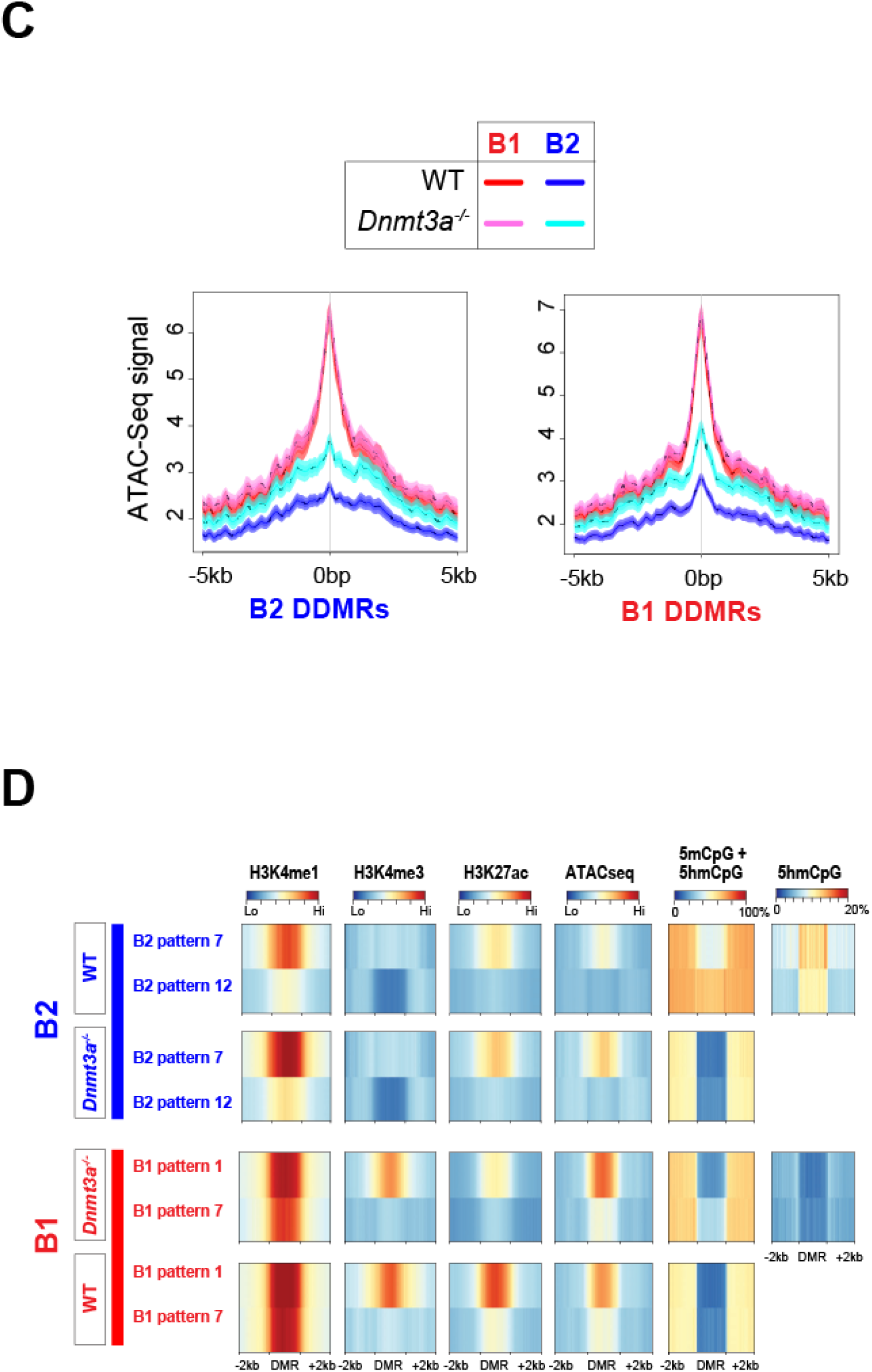
DDMRs exhibit enhancer marks that are modulated by CpG modification patterns. (A and B) 5mCpG, H3K4me1, H3K27Ac, H3K4me3 and ATAC-Seq signal in B2 and B1 DDMRs in proB, WT B and *Dnmt3a*^−/−^ B cells in the B1 and B2 lineages, respectively. Average CpG methylation ratio within the B1 and B2 DMRs is depicted alongside a heatmap of histone modifications, as measured by CUT&RUN or chromatin accessibility signals in 100 bp bins in ±5 kb regions from the center of the DMRs. The six main patterns of methylation are labeled. (C) Profiles of chromatin accessibility in B1 and B2 DDMRs as measured by ATAC-seq. (D) The average profile of modified CpGs, H3K4me1, H3K4me3, H3K27ac, and chromatin accessibility in the two largest patterns of B1 and B2 DDMRs in WT and *Dnmt3a*-deficient B1 and B2 B cells, respectively.

In B2 B cells, as is typical of active enhancers, DMEs exhibited both 5mCpG and 5hmCpG, and were marked by H3K4me1 but not H3K4me3 **(Figure 6A)**. The few B2 DMEs (pattern 1) that were enriched in H3K4me3 were also enriched in TSS, suggesting that they are promoters **(Figure S5)**. In contrast, in B1a B cells, the developmentally hypomethylated DMEs (especially pattern 1) exhibited both H3K4me1 and H3K4me3 marks **(Figure 6B)**, although they exhibited a minimal overlap with promoters **(Figure 6D and S5)**. As illustrated in **Figure 2C** using publicly available ChIP-seq data, and in **Figure 6B** with a different antibody in a CUT&RUN assay, we show that ‘promoterization’ of DMEs is a unique feature of B1a B cell development. Although CpG methylation in the DMEs was largely erased in the Dnmt3a-deficient B1a and B2 B cells, the ablation of *Dnmt3a* had little impact on H3K4me1 and H3K4me3 levels in the DMEs in either B1a or B2 B cells (**Figure 6A and 6B**). Thus, the loss of CpG methylation did not result in the acquisition of non-canonical enhancer marks by B2 B cells. Complete loss of CpG methylation in the DMEs in *Dnmt3a*-deficient B1 B cells was also associated with an increase in H3K27ac marks in the B1 DMEs bearing non-canonical H3K4me3 enhancer marks (Pattern 1), suggesting that CpG methylation may constrain the activity of enhancers with non-canonical H3K4me3 marks **(Figure 6B and 6D)**. In contrast, in B2 B cells, the levels of H3K27ac, H3K4me1 or H3K4me3 in the DMEs were largely unchanged upon DNMT3A loss.

We also observed that developmentally hypermethylated B1 DDMRs (Patterns 4 and 9) bore the H3K4me1 mark at the proB1 stage but lost it in mature B1 B cells, suggesting that these DDMRs represent enhancers that were developmentally decommissioned by DNMT3A-mediated CpG methylation. Failure to decommission this set of proB1 enhancers in *Dnmt3a*-deficient B1 B cells may have contributed to the preleukemic phenotype observed in *Dnmt3a*-deficient B1 B cells. The developmentally methylated B1 DDMRs also do not exhibit 5hmCpG marks, suggesting that DNMT3A may not be required for their continued methylation. Failure to methylate patterns 4 and 9 DDMRs is not accompanied by H3K4me3 acquisition. The impact of DNMT3A loss on the pattern of histone marks and cytosine modifications in the two largest categories of B1 and B2 DDMRs is depicted in Figure 5D.

### Dnmt3a-dependent CpG methylation of DMEs controls lineage-specific gene expression in B cells

The global loss of DME methylation in *Dnmt3a*-deficient B cells resulted in widespread perturbation in gene expression in both B1 (2,872 genes) and B2 (2,018 genes) B cells. The expression of 1,002 of these genes was concordantly altered in B1 and B2 B cells, and the expression of 1,710 and 946 genes was altered in a lineage-specific manner in B1a and B2 B cells, respectively (**Figure 7A and 7B**). In both B1 and B2 B cells, there was little correlation between the differential expression of genes upon DNMT3A loss and the presence of a DME in their vicinity (**Figure 7C**). However, DNMT3A deficiency specifically impacted the expression of B1 and B2 lineage-specific genes as well as genes that are altered during B cell development in both B1 and B2 B cells (**Figure 7C**). In both *Dnmt3a*-deficient B1 and B2 B cells, the complete loss of CpG methylation at the DMEs led to a further increase in the expression of developmentally upregulated genes in the B1 and B2 lineages. We had previously noticed that the gene closest to nearly one-third of the hypomethylation canyons in Dnmt3a−/−B cells encoded a transcription factor (TF). This suggested to us that although the presence of DMEs was not directly correlated with global changes in gene expression upon DNMT3A loss, enhancer demethylation at TF genes may result in widespread secondary changes in gene expression. Indeed, we found that the TF genes that were upregulated in *Dnmt3a*-deficient B cells were selectively enriched in the vicinity of DMEs in both B1 and B2 B cells **(Figure 7C)**.

**Figure 7:**
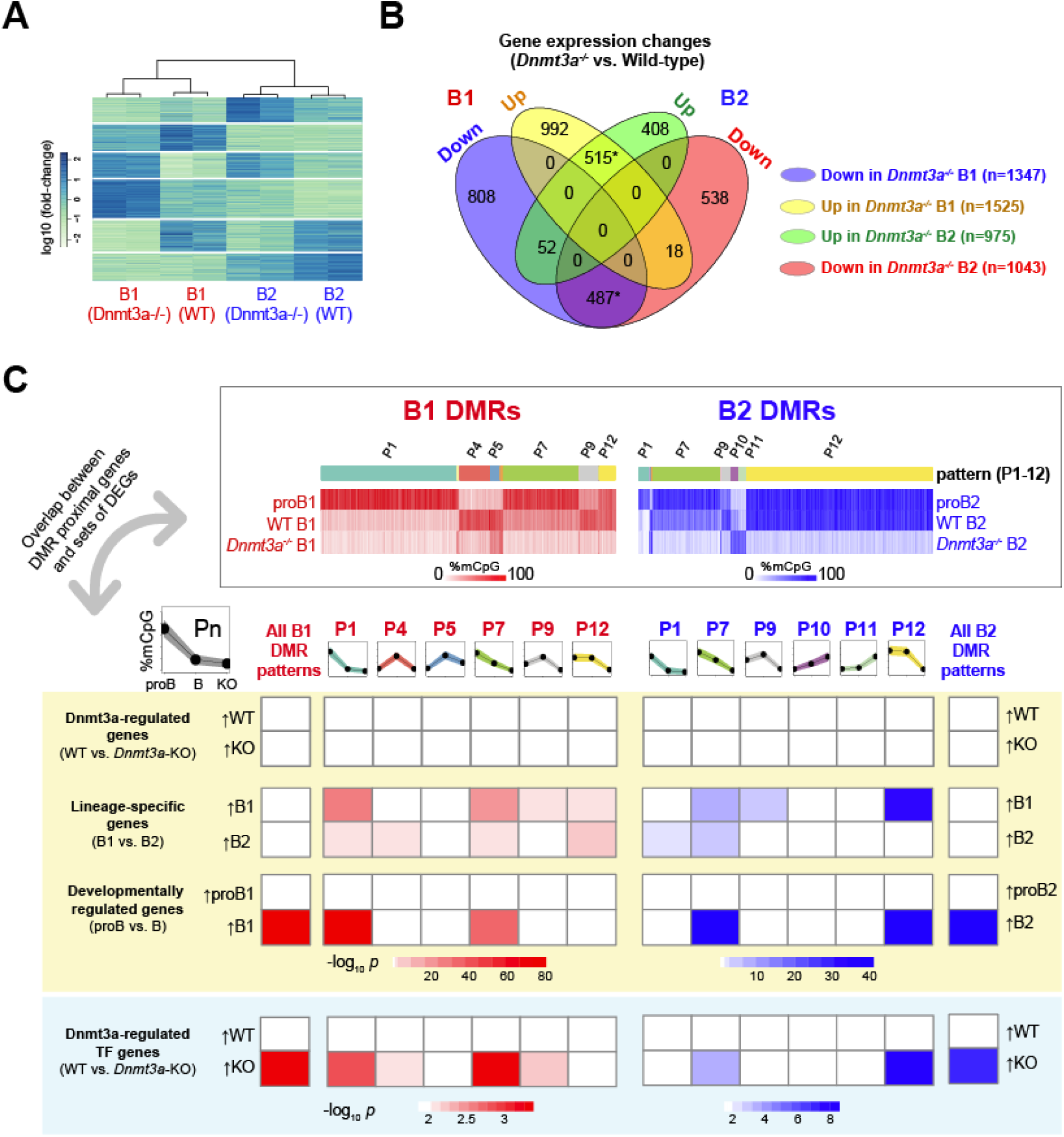
B-lineage DMRs are maintained in distinct chromatin states in B2 and B1a B cells. (A) Heatmap of differentially expressed genes between wild-type and *Dnmt3a*^*−/−*^ B1 and B2 B cells. (B) Genes differentially expressed between wild-type and *Dnmt3a*^*-/*-^ B1 and B2 B cells are shown as a Venn diagram. Statistically significant overlaps (p < 0.01, Fisher’s exact test) are marked with an asterisk. (C) Significance of overlap between genes in the proximity of B1a and B2 DMRs exhibiting distinct patterns of methylation and (i) all genes whose expression is altered by the loss of *Dnmt3a* in B1a and B2 cells, or (ii) genes whose expression is altered during development (from proB to mature B), or (iii) B2 and B1a lineage-specific genes, or (iv) only the subset of genes encoding transcription factors whose expression is altered by the loss of *Dnmt3a*. Numbers in the table are -log10(p-values) (Fisher’s exact test), and significantly overlapping pairs of gene sets (p < 0.001) are colored in pink (B1) or blue (B2) based on the p-value.

During development, lineage-specific enhancers arise at sites of silent chromatin progressing through “primed”, or “poised” states to the “active” state, each state being marked by specific combinations of chromatin and cytosine modifications ^41^. These transitions require site-specific recruitment of chromatin and cytosine-modifying enzymes through the action of TFs. We analyzed the enrichment of TF binding motifs in the DMEs and interpreted the data in the context of cell-stage specific methylation states. Indeed, DMEs with similar patterns of methylation exhibited similar TF motif enrichment, suggesting that particular sets of TFs may be responsible for specific methylation patterns **(Figure 8A)**. In any given B cell developmental stage, the CpG methylation state of DMEs is likely determined by TF-mediated site-specific recruitment of DNMT3A or TET enzymes ^42–44^. Binding motifs for TFs that have been shown to recruit to DNMT3A such as SPI1 (PU.1), FOS, JUN, and ETS1 ^42^, as well as motifs for TET2-interacting TFs such as EBF1, BATF, TCF3 (E2A), SPI1 and EBF1, were both strongly enriched in the DMEs ^43,44^. The transcriptional activator domains of the listed TFs likely determine the enhancer function of the DMEs, and it is noteworthy that the enriched TF binding motifs include both well-known master regulators of the B-lineage as well as transcriptional activators in response to BCR signaling.

**Figure 8:**
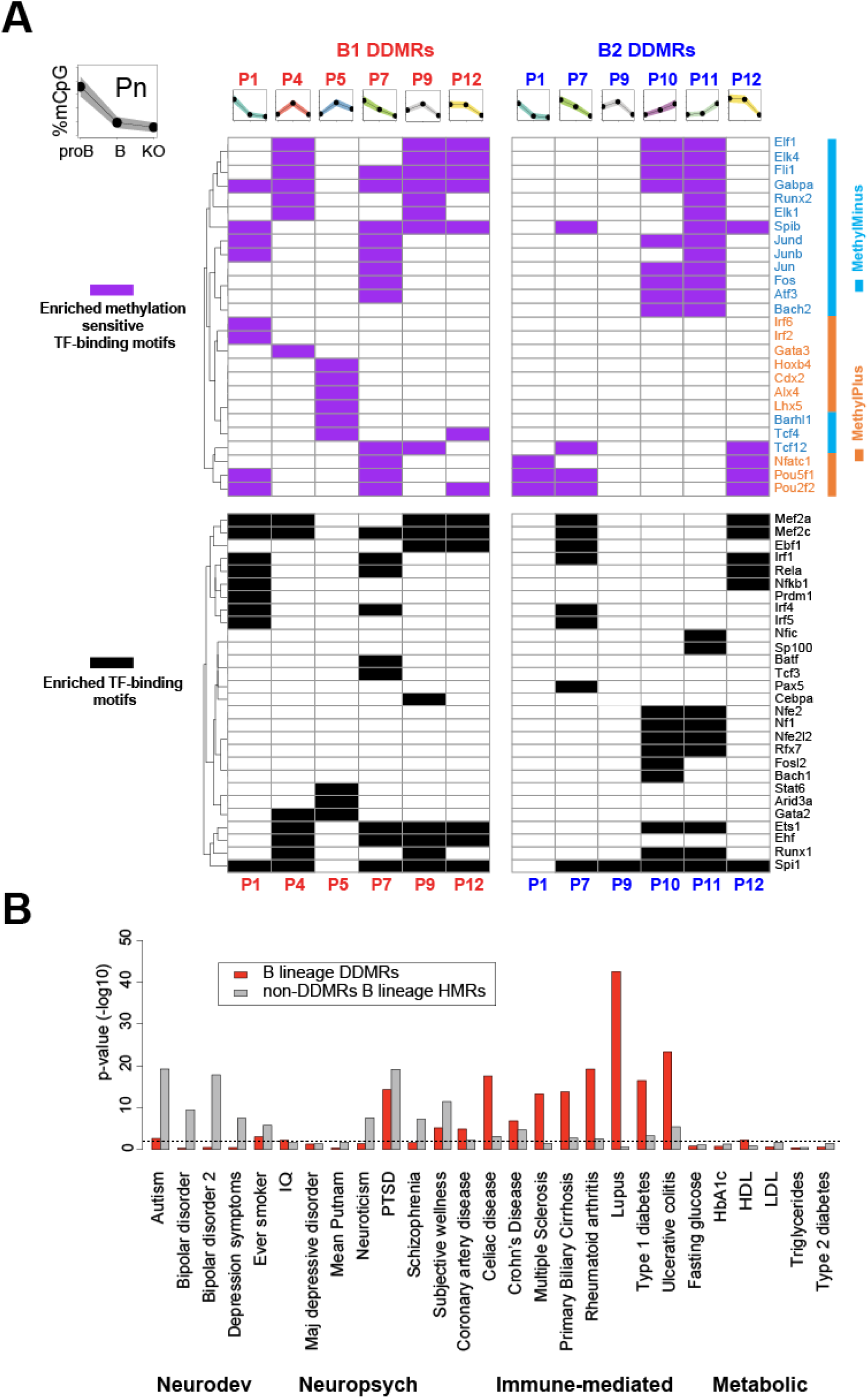
DMEs are enriched for TF binding motifs and SNPs associated with immune-mediated diseases. (A) Significantly enriched TF binding motifs in B1 and B2 DDMRs with varying patterns of methylation. Enriched motifs were filtered by p < 0.01 (binomial test run with HOMER) and a difference in motif instances between test and background sets of intervals greater than 5%; the entire set of DDMRs was used as the background. Motifs that are insensitive to CpG methylation or lack a CpG are depicted in black. Methylation-sensitive transcription factor binding motifs are marked in magenta and colored based on whether the colored by TF binding is enhanced (orange) or inhibited (cyan) by CpG methylation. (B) Enrichment of SNPs linked to human disease in GWAS in B lineage DMRs (red) and all HMRs excluding the DMRs (grey) calculated using *RiVIERA*.

Interestingly, TFs not only recruit writers and erasers of cytosine modifications, but some can also function as readers of CpG methylation as their binding can be influenced by the degree of CpG methylation at their recognition sites. The impact of cytosine methylation on TF binding has been experimentally determined in vitro for over 500 TF’s ^45^. About half of the TF binding motifs enriched in DMEs had been reported to be sensitive to CpG methylation in the Yin *et al.* study ^45^. Interestingly, the enriched TF-binding motifs for TFs with vital roles in B lineage specification, such EBF1, TCF3 (E2A), MEF2A, MEF2C, SP1 (PU.1), PAX5, and NFKB1 were either not sensitive to methylation or lacked a CpG site ^46^. They were enriched in both methylated and unmethylated intervals **(Figure 8A)**. ARID3A, which lacks CpGs in its binding motif and can selectively drive B1 lineage development, is selectively enriched in B1 DMEs in the pattern 5 intervals that are methylated during B1 development ^29^. In contrast, TFs that are activated in response to BCR signaling such as FOS, JUN, and NFATC1 are sensitive to CpG methylation in their binding sites. The binding of FOS and JUN to their targets is decreased by CpG methylation and the binding of NFATC1 is increased by CpG methylation. FOS and JUN motifs are enriched in hypomethylated intervals in both B1 and B2 B cells. Indeed, the developmental demethylation of DMEs in the B1 lineage may render a large number of DMEs responsive to FOS and JUN, which are downstream of BCR signaling.

These B lineage DMEs exhibit elevated sequence conservation across 60 vertebrate species relative to their flanking regions (**Figure S4B**). SNPs within the corresponding intervals in the human genome have been linked to human disease in genome-wide association studies (GWAS) studies ^47^. In particular, the DME syntenic regions were strongly enriched for SNPs associated with multiple human immune-mediated disorders (**Figure 8B**) ^48^, but a similar enrichment was not seen for most neurodevelopmental, neuropsychiatric or metabolic disorders. Common variants implicated by GWAS are known to be enriched for regulatory variants and it is indeed possible that the disease-associated SNPs in these DME-syntenic regions directly contribute to disease susceptibility by impacting their enhancer function ^49^. From a functional standpoint, the DMEs identified in this study can be considered to be a conserved set of methylation-sensitive enhancers in the B lineage, whose enhancer function is influenced and maintained by DNMT3A-mediated methylation.

## Discussion

Our study has uncovered the underlying foundational methylome that is shared among B cells and maintained, likely by DNMT1, in both B1 and B2 sub-lineages independent of DNMT3A. We also describe a dynamic methylome largely comprised of enhancers that are maintained by DNMT3A in a B1 and B2 lineage-specific manner. DNM3A is required to counteract ongoing TET activity at these enhancers. We therefore refer to these enhancers as DNMT3A-maintained enhancers or DMEs. DMEs undergo widespread programmed CpG demethylation during B1 B cell development, which coincides with the induction of the expression of lineage-specific genes in the vicinity of these enhancers. These very same enhancers remain methylated by DNMT3A in B2 B cells. However, DMEs are a site of ongoing TET activity in B2 B cells, as evidenced by 5hmCpG marks, and are completely demethylated in the absence of DNMT3A. This study shows that DNMT3A can function in a maintenance role at DME sites exhibiting ongoing cycles of TET-dependent demethylation and DNMT3A-dependent maintenance re-methylation. We also show that the developmentally demethylated DMEs acquire non-canonical promoter-like H3K4me3 modifications in B1 B cells. Given the novel biology of these enhancers, unusual chromatin features, role in and sub-lineage specification, we propose that DMEs represent a novel category of enhancers and their role in other cell types remains to be explored.

Although the underlying foundational CpG methylomes that are unmasked in *Dnmt3a*-deficient B1 and B2 B cells are virtually identical in the two sub-lineages, the cells do not lose their identity. Indeed, differences in other epigenetic marks including the non-canonical H3K4me3 marks in DMEs are also preserved despite the loss of DNMT3A. This suggests that other epigenetic features including the non-canonical H3K4me3 marks in DMEs are likely to be important in maintaining B1 or B2 sub-lineage identity. However, the loss of DME methylation is not without consequence. The loss of the DNMT3A-maintained dynamic methylome in *Dnmt3a*-deficient cells results in a perturbed transcriptional program that manifests as a preleukemic state in B1 B cells.

In B1a B cells, DNMT3A activity at DMEs may be opposed, at least in part, by the acquisition of ectopic promoter-like H3K4me3 marks - a phenomenon we have termed promoterization – thereby allowing complete demethylation by the unopposed action of TET. While some studies have described the acquisition of promoter-like H3K4me3 marks at enhancers in CD8^+^ T cells and in a tumor context, these previous studies did not evaluate them in the context of cytosine modifications ^23,24^. The well-characterized mutually inhibitory relationship between CpG methylation and H3K4me3 in mammalian cells may underlie the biochemical basis of the switch in CpG methylation and chromatin modification states of DMEs between B1 and B2 B cells. In mouse embryonic stem cells, the CFP1 subunit of the SET1 H3K4 trimethylase complex, which binds unmodified CpGs, constrains H3K4 trimethylase activity to hypomethylated promoters. In the absence of CFP1, ectopic H3K4me3 can accumulate in otherwise methylated enhancers that interact with promoters, resulting in the increased expression of their target genes ^25^. Conversely, H3K4me3, unlike H3K4me0/me1, is unable to free the catalytic site of DNMT3A from its autoinhibitory ATRX-DNMT3-DNMT3L (ADD) domain, causing local inhibition of DNMT activity in its vicinity ^50,51^. Therefore, we hypothesize that the established recruitment of the CFP1 subunit of the SET1 complex, the primary H3K4 trimethylase, to unmethylated DNA at promoters, also applies in part to unmethylated DMEs ^25,52^. Once the H3K4me3 mark is established in the DMEs perhaps in response to a transient opportunity presented by TET-mediated demethylation, H3K4me3 marks may also subsequently prevent further CpG methylation by DNMT3A. Indeed, 5hmCpG is enriched at the boundaries of the DMEs in B1a B cells. Furthermore, mice lacking TET2/3 in the B lineage completely lack B1a and marginal zone B cells ^19^. It is likely that TET enzyme mediated demethylation is necessary for the H3K4 trimethylase activity at B1 DMEs.

The importance of DMEs in the B lineage is further highlighted by the fact that they are highly enriched for binding sites of B lineage-determining TFs. The striking enrichment of SNPs linked to human immune mediated disorders within the DME syntenic regions further highlights the importance of these DMEs in maintaining cellular function, and we speculate that DMEs are likely to be of broader biological significance. Future studies using tissue-specific deletion of DNMT3A in various tissues are likely to reveal key elements of DME-mediated gene regulation in various cell types. In studies of organismal development and cell differentiation, enhancers have been identified as poised, primed, or active, depending on their histone modifications and the functional states of genes in their vicinity ^41^. We propose that a subset of enhancers are DMEs and likely also undergo a similar progression from a poised or primed to an active state. But unlike conventional enhancers, DMEs continue to exhibit localized TET activity even in the active state, which is balanced by DNMT3A. 5hmC is known to be enriched in enhancers in ES cells and is considered to mark enhancer priming in models of ES cell differentiation ^53,54^. TET2 has previously been shown to drive the generation of 5hmC at enhancers in proB cells ^44^.

Maintenance of the appropriate level of each CpG modification at DMEs may require persistent localization of DNMT3A and TET2 at these sites through interactions with chromatin and transcription factors, as observed in epidermal stem cells ^55^. CpG methylation of the majority of B-lineage DMEs is maintained by DNMT3A in B cells, but not in HSCs, suggesting that the dynamic methylome is specific to a particular cell type or lineage. Changes in the transcription factor activities that recruit DNMT3A and TET enzymes to the DMEs may underlie the loss of CpG methylation seen in B1 B cells. Given that PU.1 binds DNMT3A ^42^ and its binding motif is enriched in DMEs, it is likely to be one of the transcription factors that can recruit DNMT3A to DMEs. Interestingly, the loss of PU.1 (SPI1) in B cells has been shown to result in a switch from a B2 to a B1 B cell fate, and B1 B cells exhibit a lower level of PU.1 ^56^. The promoterization of DMEs in B1 B cells may also be a consequence of constitutive BCR signaling, which was recently demonstrated to be sufficient to reprogram B2 B cells into a B1 B cell fate ^4^. Indeed, the promoterized B1 DMEs are enriched in FOS, JUN and NFATC1 motifs, supporting the notion that BCR signaling may play a critical role in the acquisition of ectopic H3K4me3 in the DMEs. On the contrary, DNMT3A loss alone is not sufficient for H3K4me3 acquisition in the DMEs in B2 B cells or in the developmentally methylated and decommissioned enhancers in B1 B cells, suggesting that the acquisition of H3K4me3 requires a B1-specific factor, likely constitutive BCR signaling or a B1 B cell specific combination of lineage-determining factors such as LIN28, ARID3A, BHLHE41 or ZBTB32.

The local balance between DNMT3A and TET2 has also been suggested to maintain hypomethylation canyon boundaries in murine HSCs. The loss of DNMT3A in HSCs results in the expansion of canyons as well as increased self-renewal ^57^. B1 B cell development is also accompanied by a genome-wide increase in the size of HMRs and in the number of canyons observed during B1 B cell development suggesting that the reduction in DNMT3A activity during B1 B cell differentiation may not be restricted to DMEs alone but may involve a global reduction in DNMT3A activity. During development of the B1 lineage, B1 B cells spontaneously acquire a CpG methylation profile that closely resembles that seen in *Dnmt3a*-deficient B1 and B2 B cells. It has been proposed that HSCs modulate their self-renewal capacity by suppressing the level of DNMT3A via miR-29, and the increase in DNMT3A caused by the loss of miR-29a in HSCs is associated with a reduction in HSC self-renewal ^58^. We speculate that analogous mechanisms may control the levels of DNMT3A in B1 B cells, and consequently their self-renewal capacity. Indeed, the direct or indirect loss of DNMT3A in the B lineage has been shown to result in the leukemic transformation of B1 B cells ^35–37^. The reduced level of CpG methylation at DMEs in B1 B cells may be sufficient to facilitate self-renewal; complete demethylation in the absence of DNMT3A in the knockout context may lead to uncontrolled self-renewal.

Our work suggests that large subsets of enhancers, modulated by DNMT3A are likely present in many differentiated cell types and the contribution of DMEs in the dynamic methylome to the differentiation and biology of other lineages will undoubtedly be very revealing. DMEs may provide an additional mode of developmental control of enhancer function, wherein promoterization and programmed CpG demethylation facilitate additional modulation of enhancer activity, particularly in the context of lineage-specific gene expression. Our work also suggests how the cellular phenotypic consequences of DNMT3A-mediated epigenetic modifications can be constrained to a specific developmental lineage and how the loss of such epigenetic regulatory mechanisms may contribute to a cell-type specific malignancy.

## Acknowledgements

We thank Eliezer Calo for helpful discussions and Deepta Bhattacharya and Ananda Roy for comments on the manuscript. This work was supported by NIH AI110495 and Ragon Institute Strategic Funding to SP and support from NIH AI113163 to VSM.

## Author contributions

SP, VM and HM conceptualized and supervised the study. VM, HM, GY, HA, MA, SM and YT performed the experiments. VM, NS and VV performed the computational analysis. AC helped establish the *Dnmt3a*-deficient mouse colony. VM and SP wrote the manuscript.

## METHODS

### Mice

Generation of knockout mice bearing the conditional *Dnmt3a* allele, in which the exon encoding the catalytic domain of DNMT3A (exon 19) is floxxed (*Dnmt3a*^*tm3.1Enl*^, originally referred to as *Dnmt3a*^*2lox*^) has been described ^33^. The mice were a gift from Drs. Taiping Chen and En Li. *Dnmt3a*^*2lox/2lox*^ mice were backcrossed into the C57BL/6J background (Jax strain:000664) for over ten generations. They were crossed into *Cd19-Cre* mice (Jax strain:006785) to obtain *Cd19-Cre*^*+/−*^ *Dnmt3a*^*2lox/2lox*^ mice with a conditional deletion of *Dnmt3a* in B cells ^34^. These mice are referred to as *Dnmt3a*^*−/−*^ mice in the manuscript for simplicity. C57BL/6J mice were used as controls. All animals were housed at the Massachusetts General Hospital under specific pathogen-free conditions according to institutional animal care and use committee guidelines.

### Flow cytometry and isolation of B cell populations

Single cell suspensions from spleens, bone marrow, and peritoneal washes were analyzed using flow cytometry. The following antibody (from Biolegend unless otherwise specified) panels were used. (i) B2 B cell panel for spleen: APC/Cy7 anti-mouse CD19 (Clone 6D5), PerCP/Cy5.5 anti-mouse/human CD45R (B220, Clone RA3-6B2), APC anti-mouse IgM (Clone RMM-1), FITC anti-mouse IgD (Clone 11-26c.2a), PE/Cy7 anti-mouse CD93 (Clone AA4.1), PE anti-mouse CD21 (Clone 7E9), Pacific Blue anti-mouse CD23 (Clone B3B4). (ii) B1a B cell panel for peritoneal wash: APC/Cy7 anti-mouse CD19, FITC anti-mouse CD3 (Clone 17A2), PE anti-mouse IgM (Clone RMM-1), Pacific Blue anti-mouse CD11b (Clone M1/70), APC anti-mouse CD5 (Clone 53-7.3), PE/Cy7 anti-mouse CD23 (Clone B3B4). B1a B cells were also analyzed in the spleen and bone marrow of these mice by replacing anti-mouse CD11b with PerCP/Cy5.5 anti-mouse/human CD45R (B220) in this panel.

For purification of proB1 cells (Fraction BC from E16 fetal liver), fetal liver cells from a total of 14 male and female E16 mouse fetuses were pooled and stained with a cocktail of APC-Cy7-conjugated antibodies against Ly6c, Ter119, Gr-1, Mac-1 and CD3. Lineage-negative cells were enriched using anti-Cy7 magnetic beads (Miltenyi Biotech). The enriched lineage-negative cell fraction was then stained with a cocktail of antibodies comprising APC anti-mouse CD3, anti-mouse IgM, Brilliant Violet 421 anti-mouse CD19, PE/Cy7 anti-mouse CD93, PerCP/Cy5.5 anti-mouse/human CD45R (B220), Alexa Fluor488 anti-mouse CD24 (Clone 30-F1) and PE anti-mouse CD43. ProB1 (Fr-BC) cells were FACS-sorted on a BD FACSAria II (BD Biosciences) following the gating strategy used by the ImmGen Consortium and included an APC-Cy7 dump gate (Ly6c^-^ Ter119^-^ Gr-1^-^ Mac1^-^ CD3^-^ CD19^+^ IgM^lo^ CD93^+^ B220^-^ CD43^+^ CD24^+^). A total of 100,000 cells were sorted in RPMI medium with 0.1% BSA and the cell pellet was frozen down. DNA was subsequently extracted using the QiaAmp DNA Mini kit. ProB2 cells (Fr-BC cells) were enriched and isolated using a similar strategy as above from the bone marrow of 4 wk-old mice. ProB2 cells (Fr-BC) were FACS sorted following the gating strategy used by the ImmGen Consortium (Ly6c^-^ Ter119^-^ Gr-1^-^ Mac1^-^ CD3^-^ CD19^+^ IgM^lo^ CD93^+^ B220^-^ CD43^+^ CD24^lo^).

Single cell suspensions were made from spleens, bone marrow, peritoneal washes or fetal liver. RBCs were lysed with 2 mL ACK lysis buffer (Lonza). Prior to surface staining, 1 x 10^6^ cells were labelled with the LIVE/DEAD Fixable Blue Dead Cell Stain kit (Thermo Scientific) at 1:1000 dilution in 1 mL of HBSS for 30 minutes at room temperature. Fc-receptors were blocked with 2.5 μg of clone 2.4G2 (Biolegend) for 20 minutes on ice. Surface staining was performed using appropriate dilutions of antibodies in 100 μL of cell staining buffer (Biolegend, CA) for 30 min in the dark at 4°C or on ice. Flow cytometry and FACS sorting was performed on a 4-laser BD LSR-II, a 5-laser BD Fortessa, or a BD FACS Aria (BD Biosciences), as appropriate. For RNA-Seq studies, live cells were directly sorted into RLT plus buffer (Qiagen) containing 1% 2-mercaptoethanol (Sigma) and stored at -80°C until the RNA extraction. For ATAC-Seq, WGBS, and TAB-Seq, live cells were sorted into media containing 1% BSA, spun at 500×g at 4°C for 10 minutes. Cells were then processed immediately for ATAC-Seq library preparation or resuspended in DNA/RNA shield (Zymo Research) and stored at - 80°C prior to DNA extraction using the QiaAmp DNA Mini kit for WGBS or TAB-Seq.

### RNA-Seq library preparation

CD19^+^ CD23^-^ IgM^+^ CD5^+^ B1a B cells (35,000-100,000 cells) were FACS sorted from the peritoneal washes of two *Dnmt3a*^*−/−*^ mice and two littermate controls at 4 weeks of age; CD19^+^ IgD^hi^ IgM^lo^ CD21^+^ follicular B cells (100,000 cells per sample) were sorted from spleens of two *Dnmt3a*^*−/−*^ mice and two littermate controls at 8 weeks of age. RNA was isolated from the sorted cells using an RNeasy-plus Micro kit (Qiagen). RNA-Seq libraries were prepared using the KAPA Stranded RNA-Seq Library Preparation kit (KAPA Biosystems). Libraries were assessed for quality using High Sensitivity DNA chips on the Agilent Bioanalyzer, quantified using Qubit fluorometer (Thermo Fisher Scientific) as well as KAPA Library Quantification kit (KAPA Biosystems), and sequenced on the Illumina NextSeq 550 platform.

### WGBS library preparation

DNA was isolated from peritoneal cavity B1a B cells, splenic follicular B cells (*Dnmt3a*^*−/−*^ vs WT), fetal-liver proB1 cells, and bone-marrow proB2 B cells. WGBS libraries were prepared using 100 ng of DNA using the Pico Methyl-Seq™ Library Prep Kit (Zymo Research) according to the manufacturer’s protocol. The quality of the libraries was assessed using High Sensitivity D5000 Screen Tapes (Agilent 4200 Tapestation) and quantified with KAPA Library Quantification kit. Libraries were sequenced on an Illumina NextSeq 550. Fewer CpGs were modified in proB1 and B1 B cells (74.1% and 73.1%, respectively) than in the proB2 and B2 B cells (75.9% and 76%, respectively) **(Figure S1)**. Since CHH and CHG methylation (where H represents A, T or C) represented less than 2% of all cytosine methylation in proB and B cells **(Figure S1)**, we focused our analyses on CpG modifications.

### TAB-Seq library preparation

TAB-Seq libraries were constructed as described (Wisegene, catalog #K001) ^59^. Briefly, 5-hydroxymethyl cytosines in 500 ng genomic DNA with 0.1% spike-in control DNA (synthesized with 5-hydroxymethyl cytosines or 5-methyl cytosines exclusively) were protected using T4 β-glucosyltransferase, and 5-methyl cytosines were oxidized using recombinant TET1. Following TET1 oxidation, the DNA was subjected to bisulfite conversion and PCR-based sequencing adapter tagging as described for WGBS above (Pico Methyl-Seq Library Prep Kit, Zymo Research). The library was quantified and sequenced on an Illumina Nextseq 550. Genome-wide, 2.6% of CpGs were hydroxymethylated in B1 B cells and 3.8% of CpGs were hydroxymethylated in B2 B cells **(Figure S1**).

### ATAC-Seq library preparation

ATAC-Seq libraries were prepared as originally described ^40^. Fifty thousand freshly sorted cells were pelleted and washed with 50 μL chilled 1× PBS, and with 50 μL lysis buffer (10 mM Tris-HCl pH 7.4, 10 mM NaCl, 3 mM MgCl2, 0.1% IGEPAL CA-630). The nuclei were pelleted in a cold micro-centrifuge at 550×*g* for 10 min, and resuspended in a 50μL transposition reaction with 2.5μL Tn5 transposase (FC-121-1030; Illumina) to tagment open chromatin. The reaction was incubated at 37°C in a Thermomixer (Eppendorf) at 300 rpm for 30 min. Tagmented DNA was purified using a QIAGEN MinElute kit and amplified with 7 or 11 cycles of PCR, based on the results of a test qPCR. ATAC-Seq libraries were then purified using a QIAGEN PCR cleanup kit and quantified using KAPA library quantification kit (KAPA Biosystems, Roche) and sequenced on the Nextseq 550 platform.

### CUT&RUN library preparation

CUT&RUN libraries were prepared using rabbit monoclonal antibodies against H3K4me1, H3K4me3 and H3K27Ac (Cell Signaling Technology catalog numbers 5326, 9751, and 8173 respectively) as previously described ^39^. The protein-A Mnase fusion protein from a gift from Steven Henikoff.

### Preparation and analysis of Ig Heavy Chain-Repertoire libraries

RNA from B1a B cells was reverse transcribed and amplified using a Qiagen OneStep RT-PCR kit (Qiagen Inc., Valencia, CA, USA) and iRepertoire® mouse BCR heavy chain (MBHI) primers (iRepertoire Inc., Huntsville, AL, USA), following the iRepertoire user manual for the Illumina sequencing platform. Following manufacturer’s instructions in the iRepertoire manual, Illumina adapters were added using a Qiagen Multiplex PCR kit, and a 350–500 bp bands corresponding to the Ig heavy chain amplicons were gel purified using a Qiagen Gel Extraction kit. The libraries were quantified using Qubit dsDNA quant (Life Technologies) and sequenced on the Illumina MiSeq platform (paired end, 250 bp). For samples from aged mice bearing clonal leukemias, the libraries were Sanger sequenced using the Illumina universal primers as the sequencing primer. Repertoire data was analyzed using the *iRweb* online analysis platform provided by iRepertoire Inc.

### Analysis of WGBS and TAB-Seq data

The first 5 and last 2 bases were trimmed using *cutadapt v1.11* and the trimmed reads were aligned to the *mm10* mouse reference genome using *Bismark v0.14.3* ^60^ *with the parameters: --bowtie2 --non_directional --un --ambiguous --phred33-quals*. PCR duplicates were removed using *Picard* (v2.5.0). Approximately 5× coverage was obtained after removal of duplicates (Figure S1). The *Bismark* package was used to extract the coverage and methylation status of individual cytosines. We then used the *MethPipe v3.4.3* ^*20*^ to identify differentially-methylated regions. We implemented *MethPipe* in the following steps. First, we used the program *to-mr* to convert the output BAM file from *Bismark* to *mr* format. Second, we removed duplicate reads using *duplicate-remover* to get rid of the reads resulting from PCR over-amplification before calculating the methylation level. Third, the bisulfite conversion rate, defined as the rate at which unmethylated cytosines in the sample appear as thymidines in the sequenced reads, was measured by the program *bsrate* from *MethPipe* based on the non-CpG cytosines which are believed to be almost completely unmethylated in most mammalian cells. 98-99% bisulfite conversion efficiency was observed. Finally, the *methcounts* program was used to estimate the methylation level for individual cytosines, calculated as a probability based on the ratio of methylated to total mapped reads at the respective cytosines. Symmetric CpGs were merged using *symmetric-cpgs*. The *bedgraph* output from *Bismark* was converted into the *bigwig* format for visualization on *IGV* and for plotting heatmaps using the *SeqPlots* package ^61,62^. WGBS data for HSC (GSE49714) and CD8^+^ T cells (GSE107150) were additionally downloaded and reprocessed using the same methods and parameters as used in this study ^21,63^. A similar approach was used for the analysis of TAB-Seq data. Analysis of TAB-Seq spike-in controls showed an efficiency of ∼90% for TET oxidation and ∼98% for bisulfite conversion.

### Detection of HMRs (hypomethylated regions) and canyons

HMRs were called using the program *hmr* in the *MethPipe* package, which utilizes a hidden Markov model (HMM)-based approach. A length cutoff of 3.5 kb or longer was used to identified canyons, as previously described ^21^. Similarity in the HMRs among different cell types was assessed based on the Jaccard statistic using the *BEDtools* suite ^64,65^. The Jaccard statistic between the two sets of genomic intervals ranges from 0 to 1 (no overlap to complete overlap, respectively).

### Identification and annotation of DMRs

To compare two methylomes, we identified differentially methylated regions (DMRs) using the programs *methdiff* and *dmr* in the *MethPipe* package. The first step was to calculate the differential methylation score for each CpG site using *methdiff*. Then, the *methdiff* and *hmr* output files were used to compute DMRs using the program *dmr*. Finally, we obtained a filtered set of DMRs spanning at least 10 CpGs and having at least 5 significantly differentially methylated CpG as recommended by the *MethPipe* user manual. In addition, the *roimethstat* program was used to estimate the average methylation level in each DMR across all samples of interest.

To annotate the genomic features of DMRs (promoter, intergenic, intron, exon, 3′ UTR, 5′ UTR, TTS) and identify DMR-proximal genes, we used the *annotatePeaks.pl* script from *HOMER* (v4.9) ^66^. We also used the *HOMER* scripts *findGO.pl* to perform functional enrichment tests. The *findMotifsGenome.pl* script was used to identify enriched vertebrate transcription factor binding motifs from JASPAR and *HOMER* databases in the DMR sets of interest (filtered using a p-value cutoff < 0.01 and a difference between test sets and background intervals > 5%; the full set of DMRs was used as the background).

The DMRs were grouped into patterns based on the methylation dynamics across proB, mature B and *Dnmt3a*^*−/−*^ B cells. We created a list of 12 pattern vectors of interest representing CpG methylation ratios (range [0,1]) across the three conditions (low, intermediate and high) **(Figure S4A)**. In each pattern vector, “low,” “intermediate,” and “high” values were set at CpG methylation ratios of 0.2, 0.5, and 0.8, respectively. Each vector represents the CpG methylation ratios across proB, wild-type B, and *Dnmt3a*^*−/−*^ B conditions for a particular B-cell subset. We assigned each B1a and B2 DMR to the pattern vector that yielded the maximum Pearson correlation coefficient (PCC). No DMR was assigned to more than one pattern.

### ChIP-Seq and ATAC-Seq analysis

We obtained H3K4me1, H3K4me3 ChIP-Seq data in B1 and B2 cells (GSE72017) and H3K4me1 ChIP-Seq data in mouse HSCs (GSE63276) from the NCBI GEO database ^22,67^. The ATAC-Seq libraries were generated and sequenced in our laboratory as previously described ^40^. ChIP-Seq and ATAC-Seq reads were aligned to the *mm10* mouse reference genome. We removed duplicated and mitochondrial reads, and generated *bigwig* files for plotting heatmaps using the *SeqPlots* package ^62^. ATAC-Seq or ChIP-Seq peaks were called using *MACS2* with a fold change > 5 as a cutoff ^68^.

### Transcriptomic analysis

We aligned the RNA-Seq reads to the mm10 reference genome and performed transcript quantification using *RSEM v1.25.0*. Refseq gene annotations were used. We used *EBSeq* ^69^ to identify differentially expressed genes (DEGs) between two biological conditions; we used a cutoff of 0.95 for the posterior probability of differential expression reported by *EBSeq* to determine differentially expressed genes with a false discovery rate controlled at 5%. We used the *HOMER findGO.pl* script ^66^ to perform functional enrichment tests. Fisher’s exact test was used to evaluate the significance of overlap between two gene sets, using all expressed genes as the background.

We also downloaded microarray data for developing and mature B1a and B2 B cells (proB cells from bone marrow or fetal liver, follicular B cells from spleen, FrE from fetal liver or bone marrow, and B1a B cells from peritoneal cavity) from the ImmGen consortium (GSE15907) and identified differentially expressed genes (p < 0.01) using the *RankProd* R package ^28,70,71^.

### SNPs enrichment test in DMRs

We used the probabilistic framework implemented by *RiVIERA* to infer the association of DMRs and SNPs of 27 human traits from four major disease categories (NeuroDegen, NeuroPsych, Immune, and Metabolic) ^47^.

## SUPPLEMENTARY FIGURES

**Figure S1:**
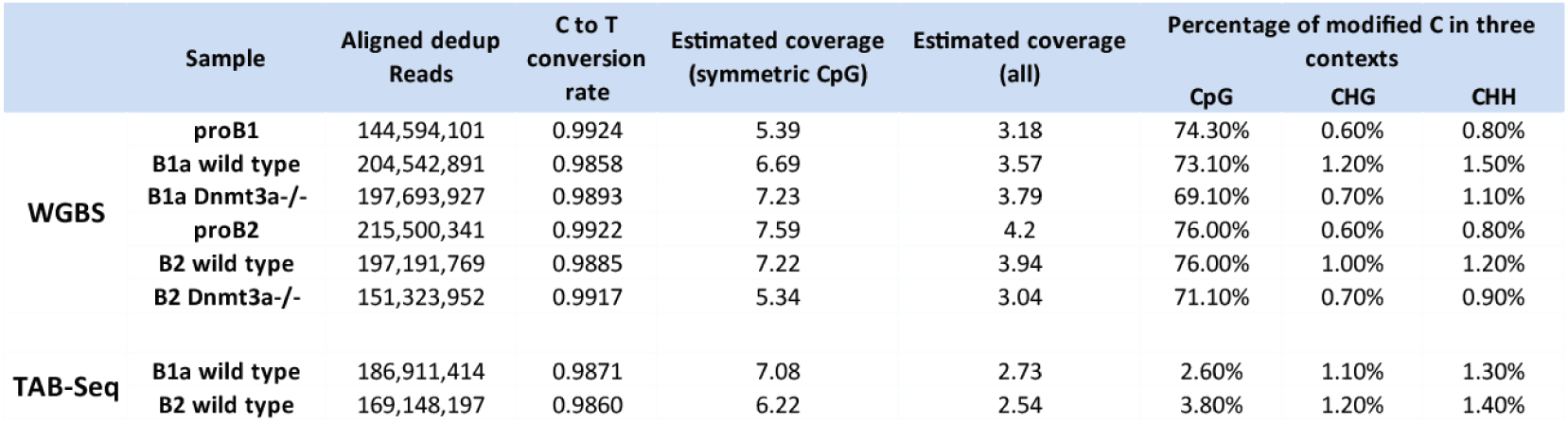
Summary of WGBS and TAB-Seq data.

**Figure S2:**
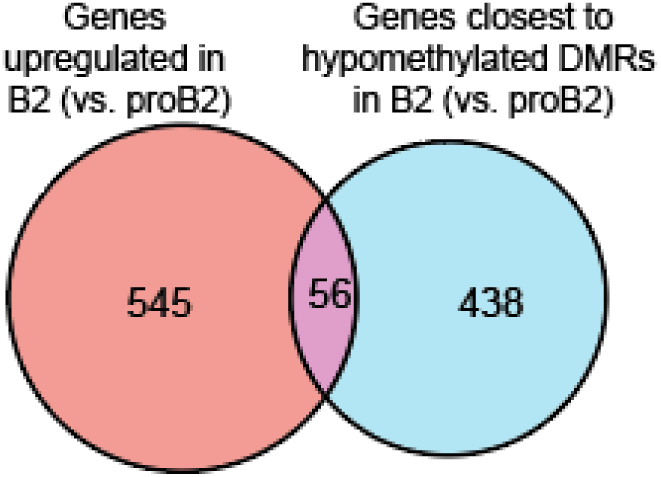
Overlap between genes upregulated during differentiation (from proB2 to B2) and genes in the proximity of hypomethylated DMRs in B2 B cells (p = 2.52e-7, Fisher’s exact test).

**Figure S3:**
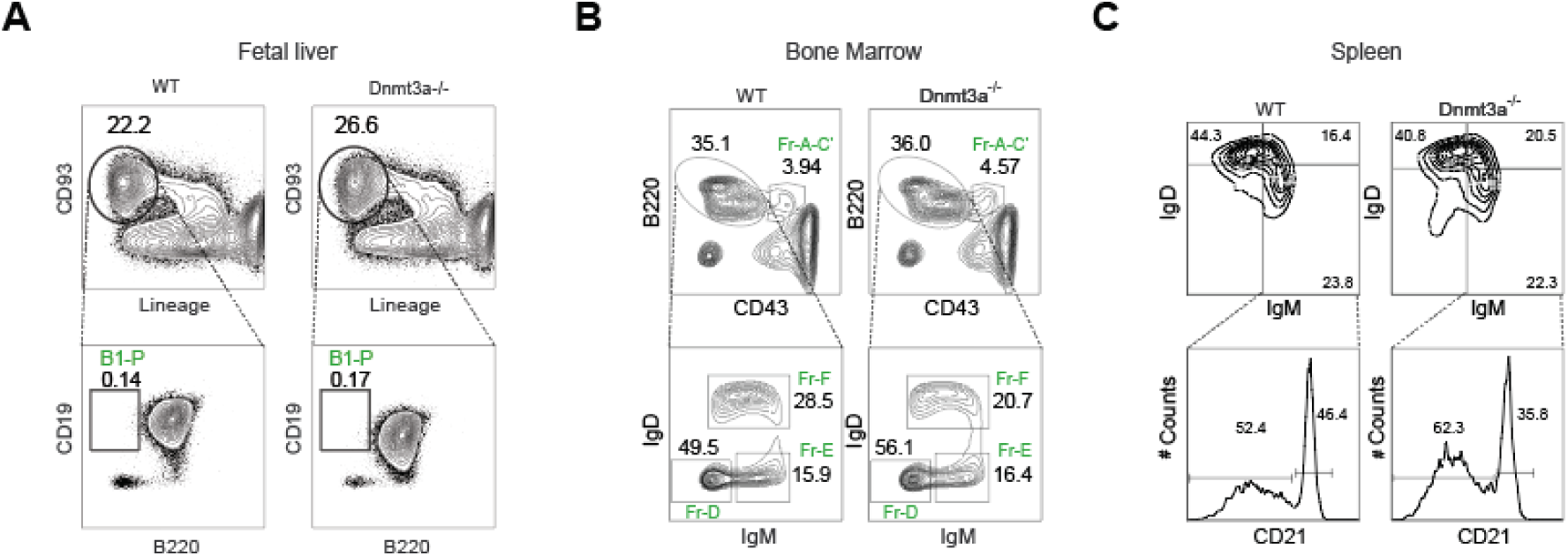
(A) Frequency of fetal liver B-1 progenitors (B1-P) in wild-type and *Dnmt3a*^*−/−*^ E16 mouse fetuses (gated as lineage-negative CD93^+^ CD19^+^ B220^-^). (B) Frequency of developing (Fractions A-E) B2 cells and recirculating (Fr-F) mature B2 subsets in the bone marrow of 4 week old wild-type and *Dnmt3a*^*−/−*^ mice. (C) Frequency of follicular (CD19^+^IgD^+^IgM^-^) and marginal zone B cells CD19^+^IgD^-^ IgM^+^CD21^+^) in the spleens of 4 week old wild-type and *Dnmt3a*^*−/−*^ mice.

**Figure S4:**
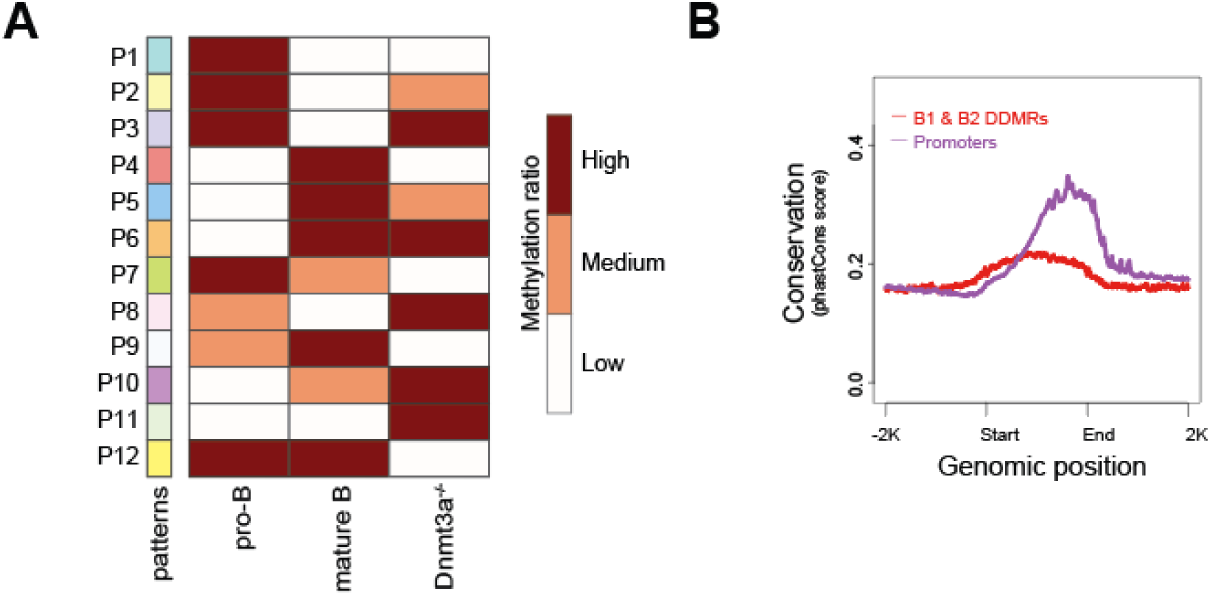
(A) Schematic of 12 possible patterns of average CpG methylation level in DDMRs across 3 cell types (proB, wild-type mature B, and *Dnmt3a*^*−/−*^ mature B cells). Each B1 and B2 DDMR was subsequently assigned to its best matched pattern. (B) DNA sequence conservation across 60 vertebrate species in B lineage DDMRs. All promoter intervals are shown for comparison. The phastCons scores for multiple alignments of 59 vertebrate genomes to the mouse genome from the UCSC phastCons60way track were used.

**Figure S5:**
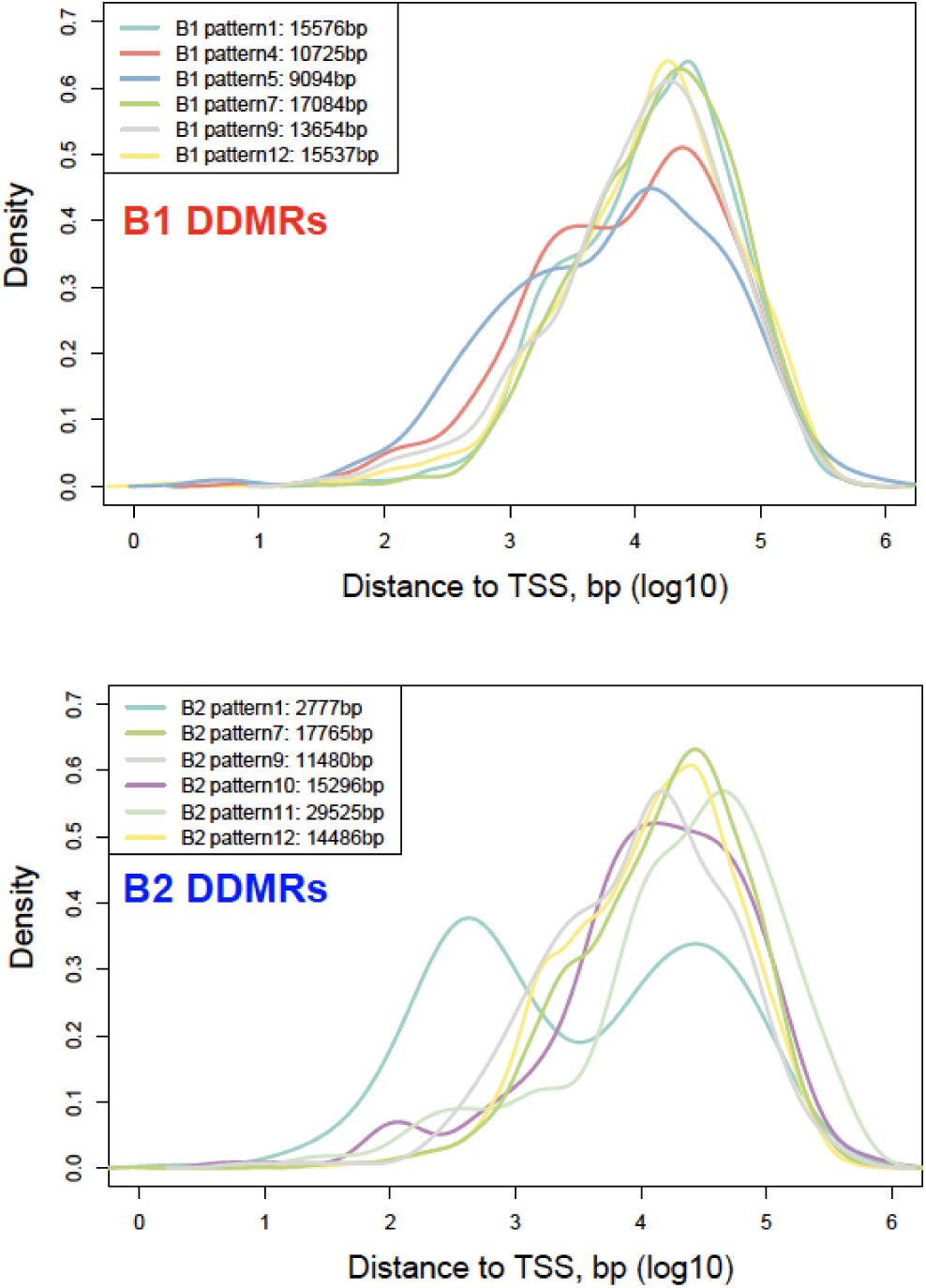
Distribution of distances between B1 or B2 DDMRs and the nearest TSS. The six most abundant DDMR patterns are depicted.

